# Tip growth in the brown alga *Ectocarpus* is controlled by a RHO-GAP-BAR domain protein independently from F-actin organisation

**DOI:** 10.1101/2021.08.28.458042

**Authors:** Zofia Nehr, Sabine Chenivesse, Bernard Billoud, Sabine Genicot, Nathalie Desban, Ioannis Theodorou, Adeel Nasir, Aude Le Bail, Hervé Rabillé, Olivier Godfroy, Christos Katsaros, Bénédicte Charrier

## Abstract

The brown alga *Ectocarpus* is a filamentous seaweed that grows by tip growth and branching. In the morphometric mutant *etoile*, tip growth is slower than in the WT and eventually stops. In this paper, we show that the causal *etoile* mutation is a null mutation in a bi-domain BAR-RhoGAP gene. By quantitative RT-PCR, we showed that *ETOILE* is ubiquitously expressed in prostrate filaments of the *Ectocarpus* sporophyte, and is downregulated in the *etoile* mutant. We immunolocalised both domains of the protein in WT and *etoile*, as well as RAC1, the known target of Rho-GAP enzymes. Thus, ETOILE would be localised at the apical cell dome where it would control the localisation of EsRAC1 to the plasma membrane. Actin staining showed that the mutant is not affected in F-actin structures. Overall, these results suggest that in Ectocarpus, BAR-RhoGAP controls tip growth by controlling RAC1 localization and through an actin-independent mechanism.

## Introduction

Tip growth is a mode of growth shared by many eukaryotic groups. The rare evo-devo study of tip-growth spanning different taxonomic taxa leads to the scenario that tip growth would derive from an amoeboid-like locomotion mechanism active in the Last Eukaryotic Common Ancestor (Vaškovičová et al., 2013), implying repeated transitions from the cytoskeleton to the cell wall as the main mechanical directional factors. Thus, in non-walled cells like amoebas and animal neurons or fibroblasts (e.g of pseudopodia, filopodia and lamellipodia respectively), tip growth relies on protruding forces generated by thick cortical bundles of parallel actin filaments (AFs) polymerizing in the front direction (Rottner and Schaks, 2019). In giant amoebas, a gradient of hydrostatic pressure generated by acto-myosin contractile forces promotes rear-to-front locomotion, therefore providing an additional protrusion force to AFs *per se*. In contrast to these “naked” cells, land plant, algal and fungal cells have all a cell wall resisting cytoskeleton mechanical forces and hence, a high turgor is required (Ali and Traas, 2016) and role of AFs is then relegated to vesicle delivery (Zhang et al., 2018).

In the filamentous brown alga *Ectocarpus*, we showed that tip growth is controlled by turgor and the thickness of the cell wall (Rabillé et al., 2019a). Put in a biophysical context, a gradient of cell wall thickness as observed in *Ectocarpus* apical cell results in a gradient of wall stress along the main cell axis. Stress is very high at the tip where the cell wall is extremely thin (∼ 36 nm) and decreases in the shanks where the cell wall is thicker (∼ 500 nm) and where growth stops. Knowing whether AFs play a role in the control of the cell wall thickness, or more generally in tip growth of *Ectocarpus* filaments, would bring an additional piece to the puzzle. Cell wall in brown algae is less stiff than in plants. It is composed of 80% of matrix polysaccharides for only ∼ 8% cellulose (reviewed in Charrier et al. (2019)). In vegetative cells, AFs are present in cortical position, where they might control the orientation of the cellulose microfibrils (Katsaros et al., 2006, 2002) similarly to the cortical microtubules (MT) in land plants, in a mode facilitating tip growth and preventing increase of thallus diameter (Paradez et al., 2006). In the greminating zygote of Fucales, F-actin was detected as patches laying in cortical position at the site of the emergence of the rhizoid prior to the germination of the zygote (Hable, 2014). There, they would initiate, through calcium gradient and MT nucleation, the accumulation of vesicles necessary for growth, from the endoplasmic reticulum to the future rhizoid pole. Thus, F-actin was shown to be necessary for the establishment of cell polarisation of the zygote of Fucales (Hable, 2014). Later during germination, F-actin organises as a ring structure in the dome of the germinating rhizoid apex.

The different roles played by actin as described above, were inferred from both pattern of labile AFs chemically labelled in fixed cells, and of morphological responses of living algae to actin depolymerising drugs. Both are prone to artefacts, that are major potential flaws in the interpreation of the data (Hable, 2014; Katsaros et al., 2006). Therefore, to know more about the functional role of AF in tip growth of brown algal vegetative cells, we first characterised genetically the *Ectocarpus* mutant *etoile* that was previously shown to be impaired in tip growth (Le Bail et al., 2011; Fig 1A) and studied the organisation of AFs in this mutant background. In this mutant, when apical cells emerge from germination or branching, they first elongate and then rapidly stop growth (Fig 1B). Here, we show that ETOILE is a RHO-GAP protein involved in tip growth. Our results further suggest that ETOILE positively regulate Rho-GTPase RAC1 activity by controlling its localisation in the tip of the apical cells. All this independently from the formation of AFs.

**Figure 1.**
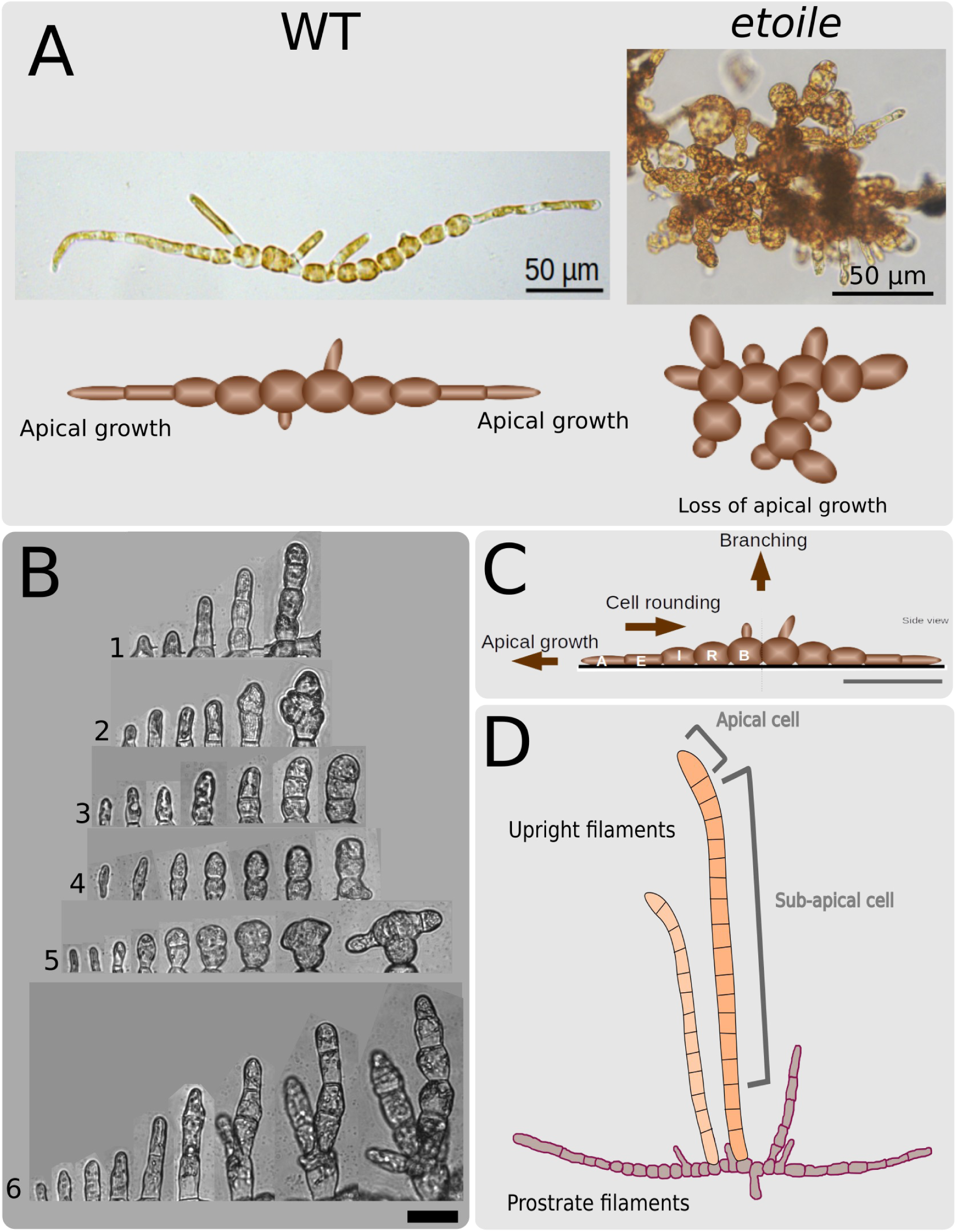
The Ectocarpus mutant etoile is impaired in tip growth. **A**. Schematic representation of early stage WT and *etoile* (*etl*) sporophytes are shown below the photos. In e*tl*, apical cells are bulky with a determinate growth, in contrast to the WT apical cells, which have an indeterminate growth mode. **B**. Time-lapse series (1-6) of growing *etl* apical cells. Cells first elongate, then stop growing and divide. This results in chunky, short cells. Note that the rounding process is not inhibited. Photos were taken every 12 hours. Scale bar 20µm. **C**. Cell differentiation in the early prostrate filament of the sporophyte. Cell rounding proceeds from apical cell elongation and division. Branching takes place on all cells except the apical cell. **D**. Cell differentiation through the emergence of upright filaments in the more mature sporophyte. These filaments grow by intercalary growth. Apical and sub-apical cells are indicated.

## Results

### The causal gene *ETOILE* codes for a BAR Rho-GAP protein

The mutation *etoile* (*etl*) was identified by positional cloning based on a progeny of ∼ 800 descendants of a cross Ec568 × etl (genetic backgroun Ec32), that were screened for the presence of Ec568/Ec32 polymorphic microsatellite markers (detailed process in Methods). Correct assembly of the genomic sequences (Suppl Fig 1 and Fig 2A) led to a predicted gene *ETOILE* (*ETL*) of 14kb comprising 12 exons (Fig 2B). Its coding sequence is long of 1740 nt and is translated into a polypeptide of 579 AA. The protein ETOILE (ETL) is composed of two domains identified by InterProscan: one AH/BAR domain (IPR027267) and one Rho-GAP domain (IPR000198 and IPR008936). The BAR domain of ETL was characterised by comparison with other BAR domains found in plants, animals and fungi (Fig 3A). Results show that the sequence of the ETL BAR domain was more similar to that of the fungal BAR domains than those of animals and plants. In particular, a phylogenetic analysis shows that ETL BAR clusters with *Magnaporthe oryzae* N-BAR domain. However, it is most likely not a true N-BAR domain, because its first amino acid is positioned 103 AA downstream from the start codon, while the H0 amphipathic helix preceding the N-BAR domain and anchoring it to the membrane is usually only ∼ 20 AA long (Peter et al., 2004). In addition, no significant portion of the leading 100 AA exhibits a clear amphipatic structure. The Rho-GAP domain shows significant conservation with known Rho-GAP proteins. In particular, alignment of the sequence with a typical vertebrate Rho-GAP (Fig 3B) identifies conserved residues, including those known to form the catalytic site of the enzyme. Each of the two domains, BAR and Rho-GAP, has been found in other *Ectocarpus* proteins (Fig 3C,D), but their combination as observed in ETL is unique in the known proteome.

**Figure 2.**
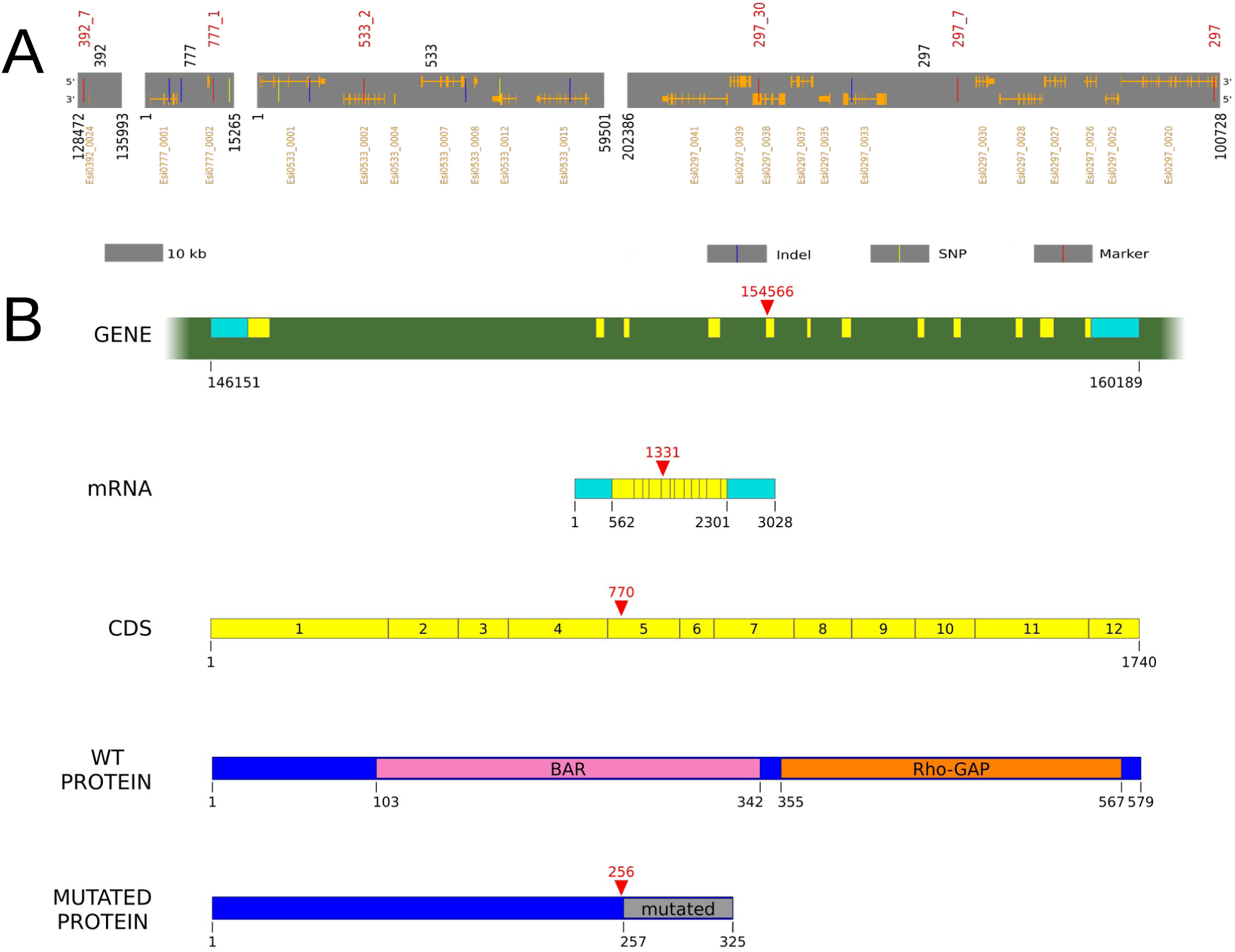
Organisation of the ETL locus, gene, coding and protein sequences. **A**. Map of the locus containing the gene *ETOILE*. The locus spans over ∼ 200kb and contains 22 identified genes (in orange, thick bars denote exons). Genetic markers are shown in red and polymorphism between the WT and *etl* is shown in blue (indels) and yellow (SNPs). **B**. Map of the gene *ETOILE* and the protein ETOILE. From top to bottom: positions of the exons on the pseudo-chromosomes (after we reconstructed it to fix the erroneously oriented super-contig 533), yellow = protein-coding, sky-blue = UTRs, red arrowhead: mutation causing the *etoile* phenotype; mature mRNA (same color code); coding sequence, with the exons numbered; wild-type protein, with the two domains identified; protein resulting from the frameshit mutation, causing changes in the AA sequence and premature translation STOP.

**Figure 3.**
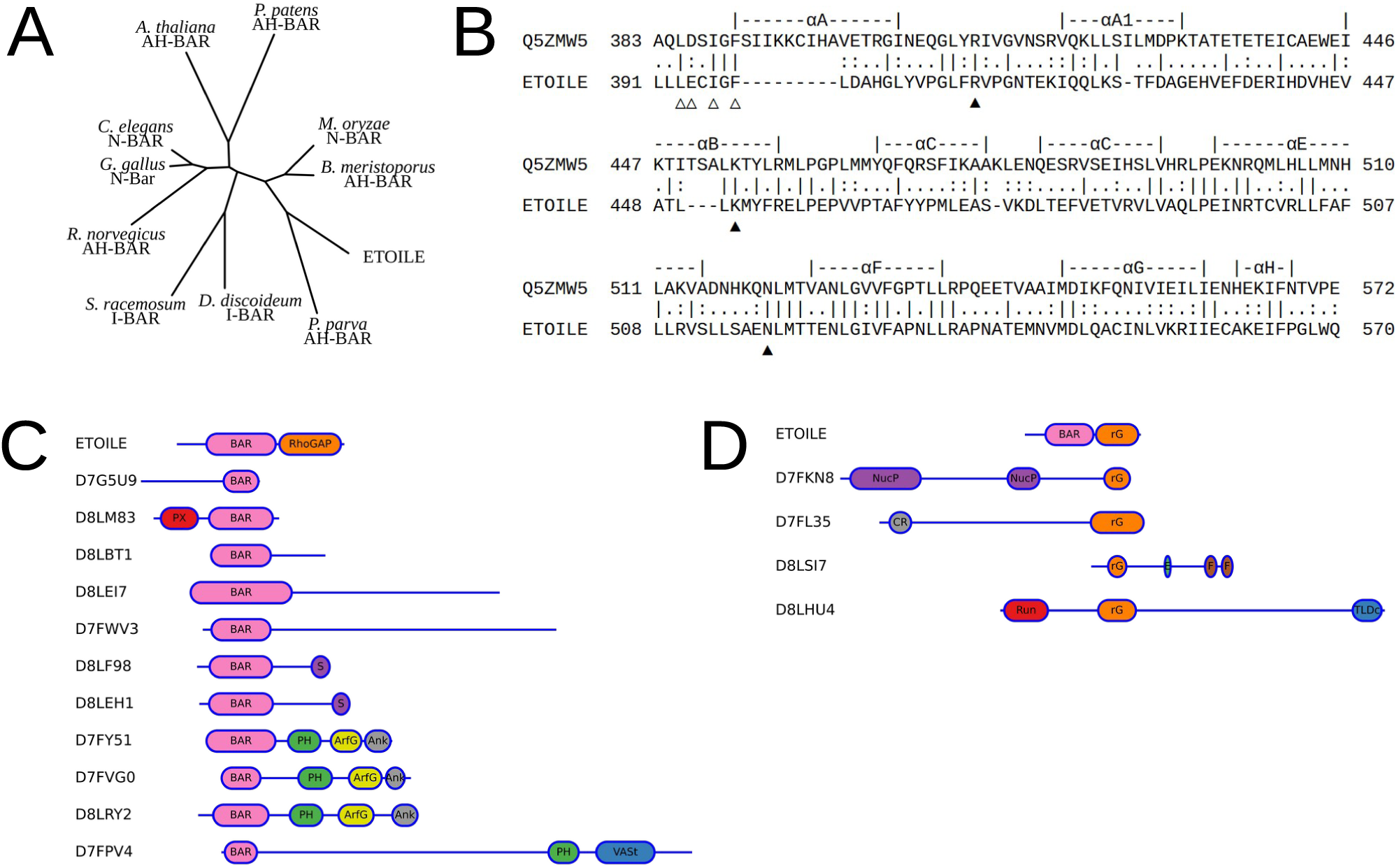
Similarity of the ETOILE protein domains in the eukaryotic lineage. **A**. Clustering of ETOILE BAR domain with BAR domains of various species (Uniprot AC in parentheses): fungi: *Basidiobolus meristosporus* (A0A1Y1ZDD2), *Magnaporthe oryzae* (A0A4P7N523), *Syncephalastrum racemosum* (A0A1X2HTP2), animals: *Caenorhabditis elegans* (B1V8A0), *Gallus gallus* (Q8AXV1), *Homo sapiens* (P53365), *Rattus norvegicus* (F1LQX4) plants: *Arabidopsis thaliana* (Q9FG38), *Physcomitrium patens* (A0A2K1JW46), amoebozoa: *Dictyostelium discoideum* (C7FZZ0), stramenopiles: *Phaeomonas parva* (A0A7S1TYS0). **B**. Alignment of the ETOILE Rho-GAP domain with the chicken Rho-GAP, displaying the secondary structures elements (above), and essential residues (Barrett et al., 1997; Longenecker et al., 2000) △: residues involved in RhoGTPase/RhoGAP interaction, ▴: conserved R, K, N residues, essential for GAP activity. *N*.*B*. for Q5ZMW5 residue numbering in Uniprot differs from those published: L193 -> L385, etc. **C**,**D**: Structural domains in ETOILE shared by other proteins in *Ectocarpus*, identified by their Uniprot AC. Domains are drawn to scale (different scales for the two sub-figures), and abbreviated as follows: PX: Phox homology, S: Sorting nexin, ArfG: Arf-GAP Ank: Ankyrin repeat, VASt: VAD1 Analog of StAR-related lipid transfer domain, rG: Rho-GAP, NucP: Endo/Exo-nuclease Phosphatase, CR: CRAL/TRIO, E: EF-hand binding site, F: FAM13. **C**. 11 other proteins have a BAR domain, usually located in the N-terminal part of the sequence. Three of these are Arf-GTPase Activating Proteins. **D**. 4 other proteins contain a Rho-GAP domain.

### ETOILE is expressed ubiquitously in the sporophyte of the WT and down-regulated in the mutant *etl*

We analysed the expression pattern of *ETL* by Q-PCR in different developmental phases and stages of the life cycle of *Ectocarpus. ETL* was similarly expressed in both the gametophyte and the parthenosporophyte (Fig 4A), whatever the developmental stage (juvenile, adult and fertile). We then tested different treatments. Two different temperatures (4°C and 30°C), 3 different osmotic conditions (H_2_0, 2M sorbitol and 2M NaCl for 1 h 30), one mechanical stress (rotative shaking), darkness for 10 days and cell wall digestion for 4 h 30 were applied to the mature sporophytes. Fig 4A shows that *ETL* response was variable, but with no significant difference between the standard conditions and the treatments (Student’s *t* test pSP juvenile *vs* all the other stage and phase: p-value > 0.285; and test pSP mix *vs* treatment: p-value > 0.140). Therefore, this first experiment shows that *ETL* expression is not highly regulated neither by the developmental stages of the alga grown in standard culture conditions, nor by the environmental conditions.

**Figure 4.**
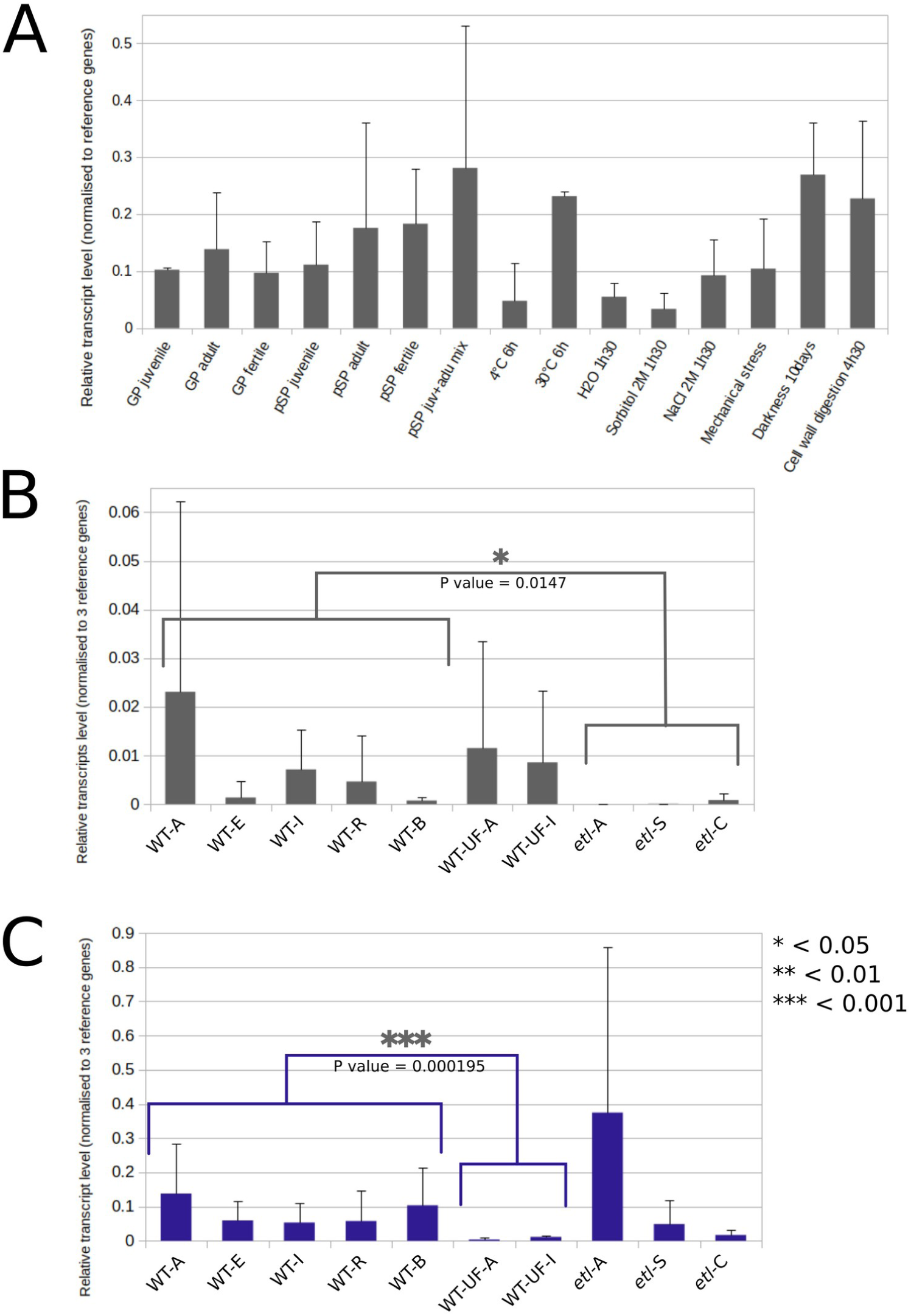
Expression of the gene etoile and EsRAC by Q-PCR. **A**. Expression level of *etl* in gametophytes (GP) at different developmental stages and parthenosporophytes (pSP) at different developmental stages and in response to different treatments. N=2 for juvenile GP and SP treatments, n=4 for all pSP. *t*-test (reciprocal and with different variance) did not show any significant differences (p-value<0.05) neither between the treatments and the pSP (mixture of juvenile and adult), nor between the pSP juvenile and all the other pSP and GP. Relative expression level is calculated as the absolute transcript number normalised to the geometric mean of the absolute transcript number of 3 reference genes (see methods). **B**. Expression level of the gene *etl* in different cell types of WT and *etl* sporophytes. *Etl* transcripts were amplified in five cell types of the prostrate filaments, namely A, E, I, R and B (see (Rabillé et al., 2019b) for the definition of the cell types) and 2 cell types of the upright filaments, namely UF-A for the apical cell and UF-I for the non-apical intercallary cells in the WT organisms. Similary, 3 cellular domain, apical (*etl*-A), sub-apical (*etl*-S) and centred (*etl*-C) domains were amplified in the mutant. n>3; *: p-value < 0.05. **C**. Expression level of the gene *EsRac1* in different cell types of WT and *etl* sporophytes. The upright filaments down-regulate the expression of *EsRac1* compared to the prostrate filaments developing earlier in the life cycle.

We then studied more deeply *ETL* expression in the sporophyte, both in the prostrate filaments and the upright filaments (also called erect, see Charrier et al. 2008). In the prostrate filament, we microdissected five different cell types, namely the apical (WT-A), the sub-apical (WT-E), the intermediate (WT-I) cells that are cells in the process of getting round, and the round cells (WT-R) present in the center of the filaments. Banching cells were also analysed (WT-B; Fig 1C). We added two cell types, the apical (WT-UF-A) and sub-apical (WT-UF-I) cells of the upright filaments (Fig 1D), that grow by intercalary growth. We performed the same microdissection in the mutant *etl*. The apical cell (*etl*-A), the sub-apical region (*etl*-S) and the central region (*etl*-C) of the prostrate base were microdissected. As up-right filaments are rare in *etl*, they were not included in this experiment. Fig 4B shows that the expression of *ETL* is ubiquitous in the WT sporophyte cell types, both prostrate and upright filaments. The high standard deviations observed in the apical cells are most likely due to the heterogeneous growing activity of these cells (some stop growing at some stage for unknown reasons and resume growth after a few hours). The primary RNA amplification due to the low amount of extracted RNA might also contribute to the overall high level of variation between biological replicates. Despite these variations, the level of *ETL* gene transcripts was significantly lower in the mutant *etl* than in the WT (p-value = 0.0147). Therefore, the D257E mutation in *etoile* might result in an increased *etl* mRNA degradation.

### ETOILE is located in the tip of the apical cell

#### *Production of antibodies against* ETOILE

In order to localise the protein *ETL* in the early *Ectocarpus* sporophytes, we generated polyclonal antibodies directed against both BAR and RHO-GAP domains separately. In a first step, both domains were overexpressed in *E. coli* and purified. The sequences chosen for each domain are shown in Suppl Fig 2. The selected areas encompass the most restrictive positions of each domain as defined by Interproscan. The overexpressed domains were used to generate polyclonal antibodies. Western blot confirmed recognition of both domains by both final sera (not shown) and the two purified antibodies (Fig 5A). When used on a total protein extract from the WT, several bands were dispayed (Fig 5B). Based on its sequence, the expected size of ETL is 62.5 kDa (Fig 5 C). A band of this size was revealed when using BAR and RHO-GAP antibodies. In *etl* and U749-51, a [*etl*] descendant of *etl* crossed twice with WT strains, these bands disappeared and new bands of ∼ 33 kDa, which is the expected size of the truncated *etl* protein (Fig 5C), were revealed by the BAR antibody. The weak signal of these bands is consistent with the very weak transcriptional pattern of the *ETL* gene as observed by Q-RT-PCR and single cell NGS analysis (not shown, refer to Billoud et al. (2021) for complete data). In addition to these weak bands, stronger bands were revealed by both antibodies. With the Rho-GAP antibodies, the strongest band was ∼ 75 kDa and weaker bands of smaller size can also be observed. With the BAR antibodies, the strongest bands are ∼ 65, 62, 50 and 40 kDa. This result indicates that the antibodies most likely recognise, in addition to *ETL*, some of the other 4 Rho-GAP domain-containing proteins and the other 10 BAR domain-containing proteins identified in the *Ectocarpus* genome. The hypothesis of cross-reactivity is supported by the fact that the genes for some of these proteins are highly expressed, as shown by quantification of their transcript levels by Q-PCR (not shown). Therefore, polyclonal antibodies are not strictly specific for ETL but most likely detect one or more of the other 14 proteins containing similar domains. This is inherent to polyclonal antibodies. Nevertheless, we relied on the co-recognition of ETL by these two antibodies to specifically immunolocalise ETL.

**Figure 5.**
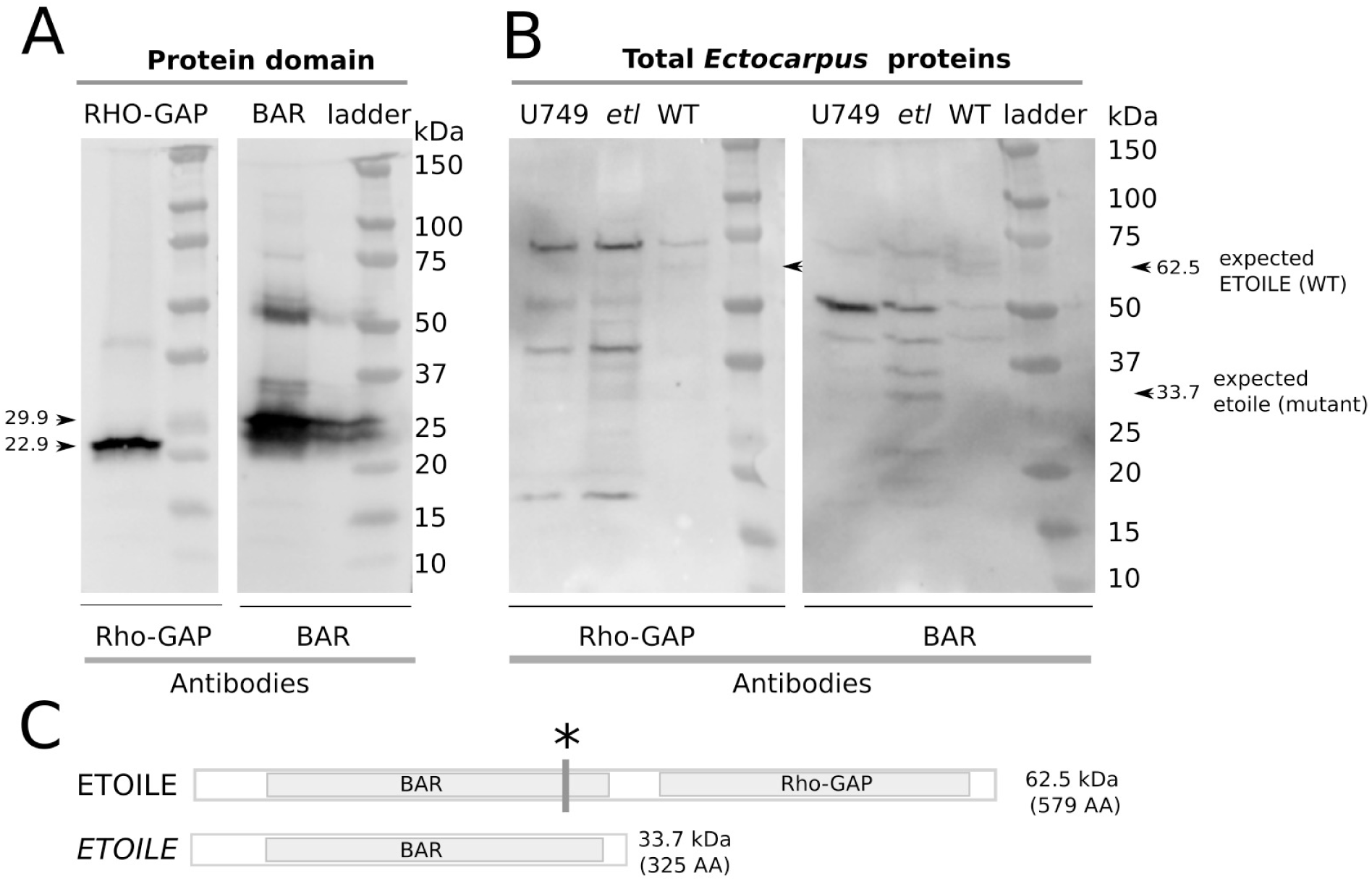
Western blot of Ectocarpus proteins with the purified polyclonal antibodies raised against ETL BAR and ETL RHO-GAP domains. **A**. Western blot run with the domains produced *in vitro*. The strongest bands correspond to the size of the BAR and Rho-GAP domains of the protein ETL, respectively 29.94 kDa and 22.99 kDa. Antibody dilution 1/1000, exposure 3 secondes. **B**. Western blot run with the *Ectocarpus* total proteins. Antibody dilution 1/1000, Exposure 10 sec. U749 is a descendant of 2 crosses of *etl* with WT. Arrows indicate the expected bands. **C**. Map of the WT ETOILE and mutant etoile proteins with expected molecular mass (kDa). Asterisk displays the location of the mutation, that makes a shift in the coding frame, leading to a stop codon ∼ 70 AA downstream of the mutation. The BAR domain is still recognised as such by Interproscan.

#### *Immunolocalisation of* ETOILE

WT filaments were immunolabelled with both ETOILE polyclonal BAR and RHO-GAP antibodies and an IgG-FITC. When used alone, this secondary antibody produced no signal (Negative control; Fig 6A-D). In apical cells, a diffuse Rho-GAP signal was detected in the most distal region of the cell, as well as some patchy signals (Fig 6E-K). Linear patches 2-3µm long located just below the dome could be clearly observed in the cortical areas (e.g. Fig 6F,G). They were either aligned with the main longitudinal axis of the cell (Fig 6E) or arranged along the curved contour of the dome area (Fig 6F,G,K).

**Figure 6.**
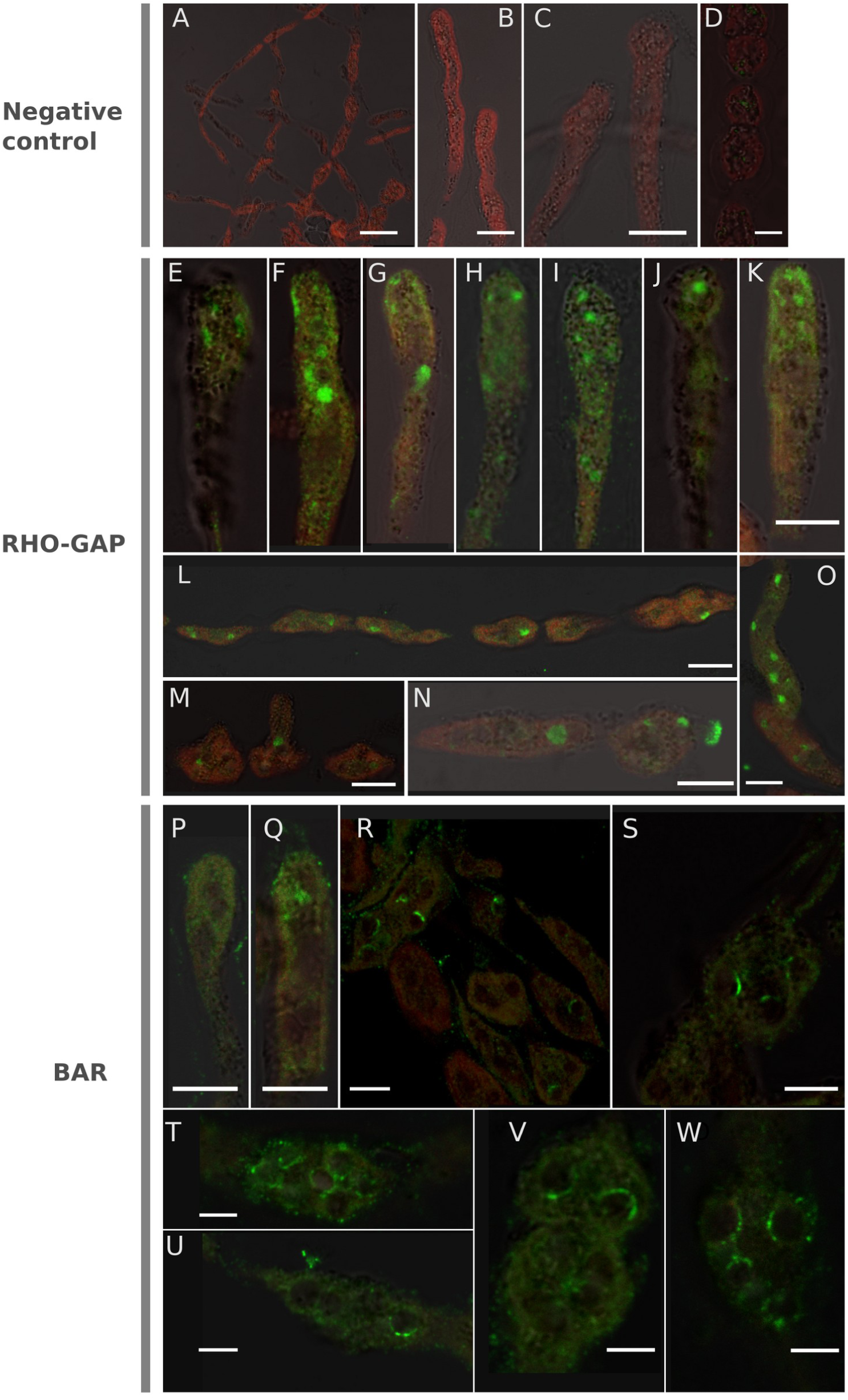
RHO-GAP and BAR antibodies recognise antigen in the apex of apical cells and in vacuoles of all cells. **A**-**D**. Negative control produced by incubation of the fixed algal sample with only the secondary antibody; **A**. overview with several filamenst; **B**,**C**. Apical cells. **D**. central, round cells. **E**-**O**. immunodetection with RHO-GAP antibody. **E**-**K**. apical cells labelled with Rho-GAP antibody. **L**-**O**. Non apical filament cells. **L**. Overview of a piece of filament, **B, N**. intermetiate, round and branching cells. **O**. germinating filament. **P**-**W**. immunodetection with the BAR antibody. **P**,**Q**. apical cell showing a diffuse signal (P) and several vesicles (Q). In contrast with the results obtained with the Rho-GAP antibody, BAR antibody labelled only the boundaries of the vacuole vesicles and not the whole vesicle. Green signal is the FITC emitted fluorescence (525nm); Red signal is the autofluorescence of the chloroplasts (> 650nm). Scale bar = 10µm in all photos except A, M and N where it is 20 µm.

In addition, a dense signal was emitted from central spots in these cells. These spots are the vacuoles, and the Rho-GAP signal covers most of its surface (Fig 6F-J). In non apical cells, these spots can also be labelled (Fig 6L-O). The BAR antibody gives a similar immunolabelling pattern (Fig 6P-W). Apical cells display a diffuse pattern mainly in the dome (Fig 6P, Q) as well as linear or dot-like signals along the dome surface (Fig 6Q). Very intriguingly, strong signals are emitted from the vacuoles, but only at their periphery, and only for a part of it (Fig 6R-W). Therefore, Rho-GAP and BAR show a similar labelling pattern. The co-immunolocalisation of these two domains, both in the dome of the apical cells, and in the central vacuoles of all cells, strongly supports that these antibodies recognised ETL. Interestingly, Rho-GAP and BAR show a slightly different labelling pattern in the vacuole: while Rho-GAP labels the entire surface (Fig 6 F-O), BAR labels only part of it, leaving the vesicle lumen unlabelled (Fig 6R-W). BAR domain usually recognises membrane curvature.

### *ETL* controls the location of EsRAC1 in the dome of the apical cells but not its expression level

The presence of a Rho-GAP domain strongly suggests that the target of ETL is a Rho-GTPase. In order to find the substrate for the Rho-GAP activity, we searched for a Rho-GTPase in the predicted proteome of *Ectocarpus*. In the UniProt database (The UniProt Consortium, 2021), one protein is annotated as “RAC, RHO family GTPase”: D8LIW3. We checked the annotation and uniqueness of this protein using profiles from the Prosite database (Sigrist et al., 2002). Within the *Ectocarpus* proteome, the PS51420 profile, described as “small GTPase Rho family profile”, corresponds to 16 proteins, including D8LIW3 and 15 other proteins, all annotated as Rab family GTPases. Note that the D8LIW3 score for PS51420 is the highest of the 16 proteins. All these 16 proteins are also matches for PS51419 (small GTPase Rab1 family profile) and PS51421 (small GTPase Ras family profile). However, for D8LIW3, the normalised scores for these three profiles are in the order Rho > Rab1 > Ras, whereas for the others 15, the order is Rab1 > Ras > Rho (See Suppl Table 3). Furthermore, the sequence of D8LIW3 alone is consistent with the sequence specificities of the Rho-GTPase family (Suppl Fig 3A), in particular the Rho-specific “insert helix” (Dvorsky and Ahmadian, 2004) and the isoprenylation site (Nguyen et al., 2010), which anchors the molecule to the membrane. We conclude that D8LIW3 is the unique Rho-GTPase protein of *Ectocarpus*.

It presents 95.9% identity and 98% similarity with the RAC1 of *Fucus distichus* (Q6T388), 71% of identity and 81% of similarity with human RAC1 (P63000) (global alignment, using the needle program of the emboss package (Rice et al., 2000)) shown to cluster with animal Rac1 by (Fowler et al., 2004). Based on this proximity and the fact that FdRac1 and the *Ectocarpus* Rho-GTPase are reciprocal best blast hit in Uniprot (e-value BLAST = 7.7.10^−137^), we named *Ectocarpus* Rho-GTPase EsRAC1. In addition we show that EsRAC1 clusters with the Racs of other stramenopiles (Phaeophycea *Fucus distichus* and oomycetes), and relates to Racs of fungi and animal, rather than other Rho family proteins, namely Rho, ROP, cdc42 (Suppl Fig 3B).

We then investigated whether EsRAC1 is expressed concomitantly with *ETL*. Figure 4C shows that EsRAC1 was expressed at similar levels in all cell types in prostrate filaments. In contrast, its expression was much lower in the erect filaments (*t* test p-value = 1.9.10^−4^). Note that the transcript level of EsRAC1 is ∼ 10 times higher than that of *ETL*. This result is very intriguing, as there is only one RhoGTPase in *Ectocarpus*, and erect filaments are an important part of the whole sporophyte thallus. However, despite their elongated shape, these upright filaments are not as polarised as the prostrate filaments, as growth is not restricted to the apex but is carried out by all cells of the filament. Therefore, this result suggests that EsRAC1 is not involved in the intercalary growth of the upright filaments.

Investigating the EsRAC1 localisation pattern, we used a commercial antibody against human RAC1 and verified its specificity against EsRAC1 by western blot. Suppl Fig 4 shows that several bands were recognised by this antibody in proteins extracted from WT and mutant algae. A band of the expected size of 21.5 kDa was displayed, but non-specific targets were also immunolabelled. Despite the presence of several bands, we were still curious as to which subcellular compartment or regions of the *Ectocarpus* cell would be labelled with the RAC1 antibody. Therefore, we performed the immunochemistry in WT and *etl* filaments and compared the pattern with that obtained with BAR ETL and Rho-GAP ETL in search of co-localised signals.

Fig 7 shows that Rac1 polyclonal antibodies significantly labelled the contour of the tip of the WT *Ectocarpus* apical cell (Fig 7A-E). In the zygote of *Silvetia compressa*, Rac1 was similarly shown to accumulate in the tip of the emerging rhizoid (Muzzy and Hable, 2013). Therefore, this similarity supports that the antibody raised against HsRac1 recognises EsRac1. HsRac1 antibody also labelled the surface of the vacuoles (Fig7 F-H). Noticeably, the signal was stronger where the curvature of the vacuoles increases during what looked like vacuole budding or fusion with smaller membrane bodies (asterisks). Interestingly, in the mutant background of *etl*, the immunolocalisation pattern was altered. It was seemingly conserved in the apical cells which kept a fairly polarised shape (the penetrance of *etl* is not complete; refer to, Le Bail et al. (2011) and Fig 1B; Fig 7I) but was delocalised in cells with a wider dome (Fig 7J-M, right). This is obvious in Fig 7M where two domes of different diameters display a different immunolocalisation pattern of EsRAC1. As in the WT, vacuoles were also labelled but with an internal signal, and not only at the surface of the tonoplast (Fig 7I, J, M). Therefore, although the Rac1 antibody can recognise several epitopes as suggested by the western blot, its localisation at the plasma membrane in the apical cell dome and its modified localisation in the context of Rho-GAP deletion strongly support that this antibody recognises EcRac1. ETL would then control the localisation of EsRAC1 at the plasma membrane. Alternatively, it is possible that in an *etl* genetic background, EsRAC1 is delocalised as the dome enlarges and therefore this mislocation is a side effect.

**Figure 7.**
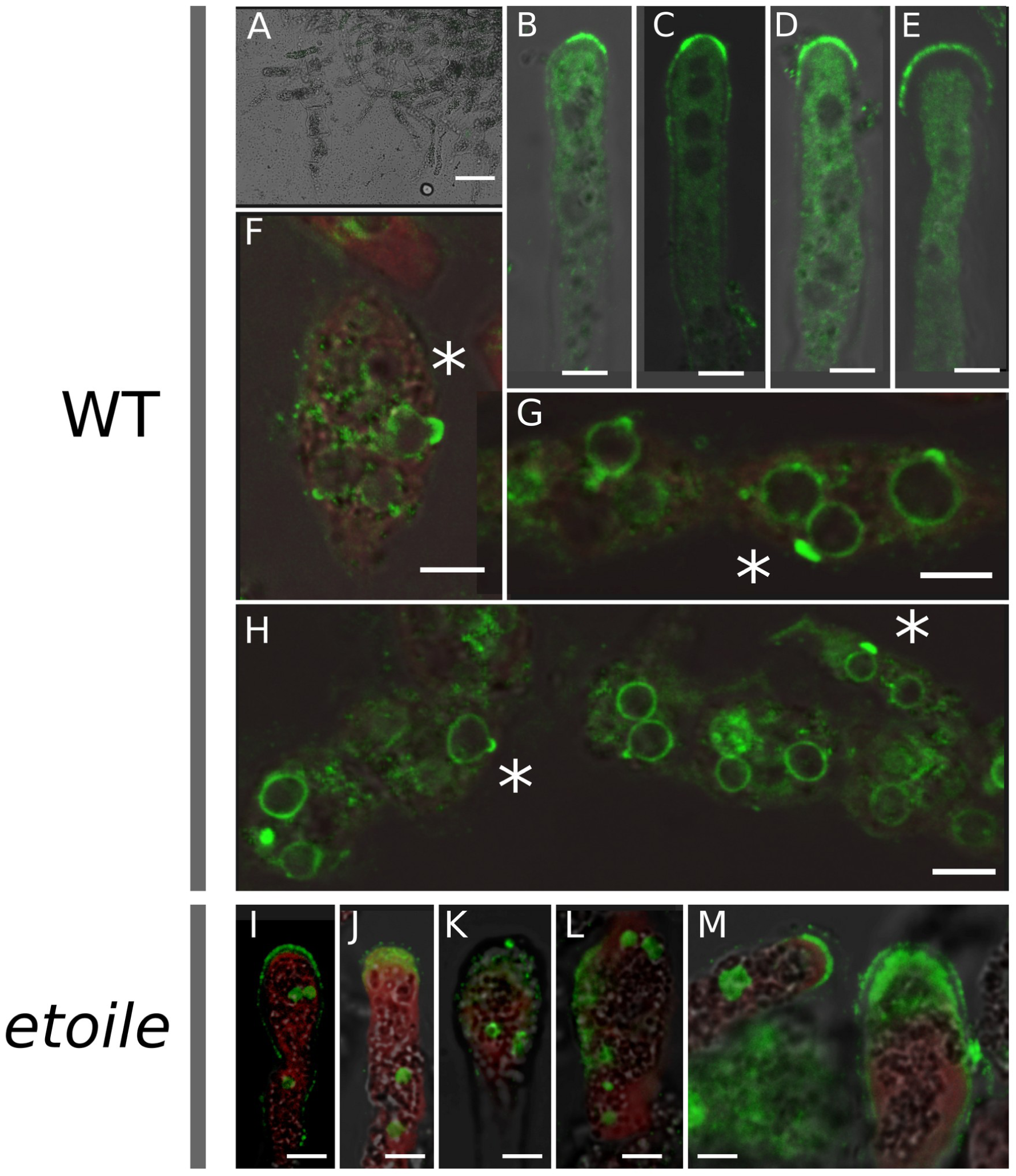
HsRac1 antibodies label epitopes in the cortex of the dome of the apical cells, and the vacuoles. HsRac1 was used with secondary antibody IgG-FITC (emision 525 nm) on WT and *etl* filaments. **A**-**H**: WT; **A**. WT negative control with only the secondary antibody showing no signal. **B**-**E**. Apical cells of WT filaments. The tip is labelled, over surfaces of different sizes. **F**-**H**. Round and intermediate cells present in the middle of WT filaments. Labelled (green) circular vesicles are vacuoles. Note the stronger signal where vacuoles are budding or fusionning (asterisk). **I**-**M**. Apical cells of *etl* filaments. The signal is diffuse in apical cells with larger diameters (K-M right). Vacuoles are also labelled. Scale bars: 25 µm (A); 5µm (B-E; I-M); 10µm (F,H).

### F-actin is not affected in *etl*

To investigate whether EsRAC1 mislocation impacts on actin organisation in *etl* apical cells, we labelled G-actin and F-actin in the WT and in the *etl* mutant. Compared to the negative control (Fig 8A), immunolabelling of G-actin with a polyclonal anti-plant actin antibody showed a diffuse but clear signal in the dome of the WT apical cell (Fig 8B,C). In *etl*, a similar pattern was observed in the apical cells (Fig 8D,E). Using Alexa 488-conjugated phalloidin, we were able to display long bundles of AFs along the apical cells of the WT (Fig 8F,G) and in sub-apical cells (Fig 8H). In the apical cells of *etl*, these bundles were also clearly displayed (Fig 8I), and even in those with a characteristically wider, round shape (Fig 8J). This is particularly convincing in Fig 8K where the AF bundles are completely intact despite the very bulky shape of this apical cell. Similarly, AF bundles were observed in the sub-apical cells of *etl* (Fig 8L). In comparison, in apical cells treated with latrunculin B, an F-actin depolymerising drug, no bundles could be observed (Fig 8M). Therefore, in *etl*, the F-actin structures are intact.

**Figure 8.**
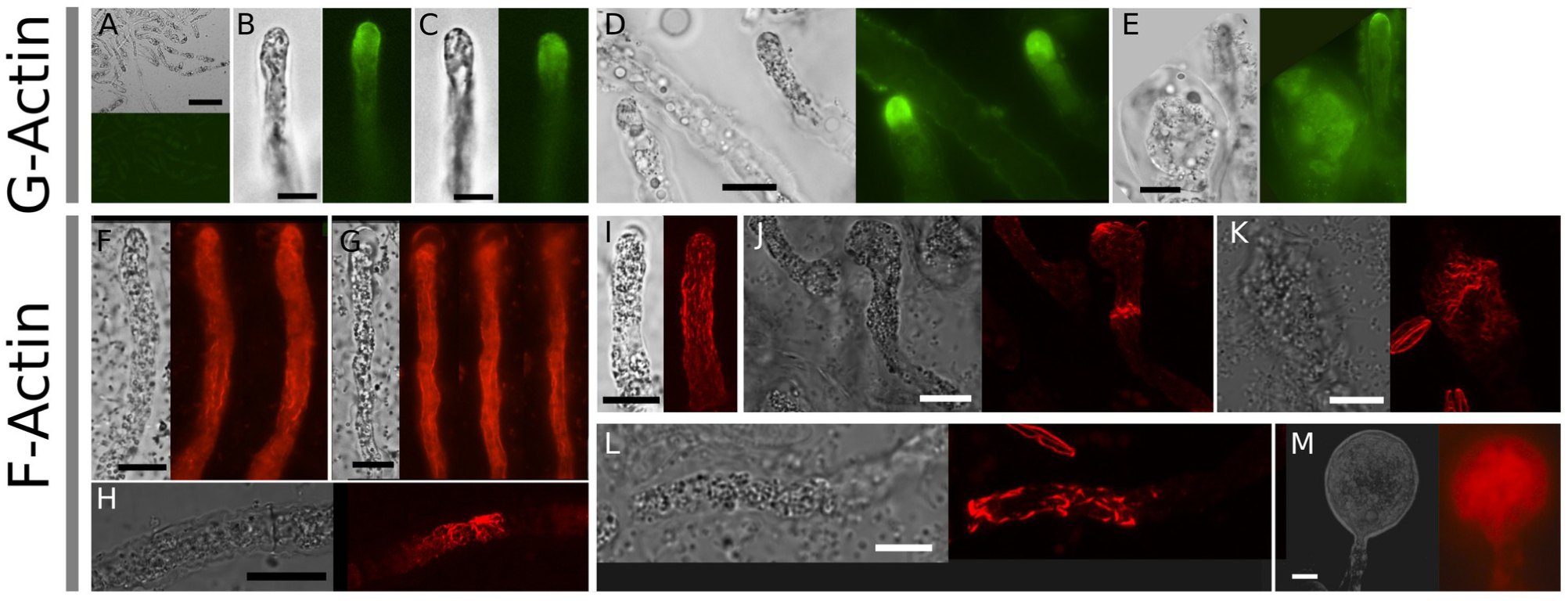
F-actin is intact in the genetic background etl. For each cell both bright field and epifluorescence or confocal microscopy are shown. **A**. Negative control for the immunolocalisation of G-actin. Only the secondary antibody was added. **B**-**E**. Immunolocalisation of G-actin with a rabbit anti-plant actin polyclonal antibody. **F**-**M**. Alexa-488-Phalloidin labelling. **B**,**C**,**F**,**G**,**H**,**M**. WT. **D**,**E**,**I**,**J**,**K**,**L**: *etl*. **M**. treated with 3µM latrunculin B in 0.1% DMSO. **A**-**G**,**M**: epifluoresence microscopy, **H**-**L**: confocal microscopy. Scale bar: 50µm in **A**, 10µm for the other photos.

## Discussion

### Rho-GAP and BAR domains in Ectocarpus

In the 1980s, the Rho-GTPase pathway was extensively studied to understand the mechanisms of polarity acquisition in the rhizoid of the Fucales embryo. AFs, the ARP2/3 complex and the Rho-GTPase Rac1 were all detected in a conical zone below the tip once the rhizoid emerged (Hable, 2014) and it was shown that reactive oxygen species (ROS) and Ca^2+^ are required for polarised rhizoid growth (Coelho et al., 2008). In 2010, these algal models were superseded by *Ectocarpus*, due to the sequencing of its genome and its suitability for genetic studies.

In this work, we identified the causal gene of the *etoile* mutant previously characterised at the phenotypic level (Le Bail et al., 2011; Nehr et al., 2011). We were unable to isolate other alleles of this mutation despite several rounds of mutagenesis from which mutants with a similar phenotype were selected. However, the fact that *etl* is a member of the Rho-GTPase pathway, which is known to be involved in cell polarisation in a wide range of organisms, leaves little doubt that we have found the right gene. Indeed, a Rho-GAP domain, present in the C-terminal region of ETL, is known to regulate Rho-GTPase activity of the Ras superfamily of small GTPases (Mosaddeghzadeh and Ahmadian, 2021) The BAR domain, present in the N-terminus, is known in other organisms to recognise membrane curvature (Peter et al., 2004). In *Ectocarpus*, 10 other proteins have a BAR domain, and 4 other proteins have a Rho-GAP domain. However, only ETL has both domains. Compared to other species, the co-occurrence of BAR and Rho-GAP is not exceptional, but it is not found in plants. More pragmatically, the presence of these two domains represented an opportunity for us to circumvent the problem of performing specific immunolocalisation of multigene family proteins. We therefore raised antibodies against each domain separately and gambled that co-immunolocating the two antibodies would show where ETL is located.

We observed a diffuse signal in the apical cell dome with both antibodies, in addition to a strong signal on the surface of vacuoles. The signal was even stronger when the curvature of the vacuole was high, suggesting that ETL is involved in, or responsive to, the dynamics of vacuole formation by budding or fusion of the tonoplast. In the etl mutated protein, the Rho-GAP domain is lost while the BAR domain should be partially preserved. As expected, the Rho-GAP antibody signal was lost in the apical cell dome (not shown). However, it was maintained on the surface of the vacuole, casting doubt on the specificity of the signal at this location. BARs and RhoGAPs other than ETL are abundant in *Ectocarpus*, both in terms of the number of genes in the genome (14 in total) and at the transcriptomic level. Therefore, a reasonable hypothesis is that some of these proteins colocalise with ETL on the surface of the vacuole. In fact, it is very likely that they overlap with its signal because *ETL* gene expression is extremely low.

### ETOILE as a topological regulator of Rac1

We have shown that in the *etl* mutant, the only Rho-GTPase enzyme encoded by *Ectocarpus* is delocalised. While in the apical cell of the prostrate filament EsRAC1 is located cortically in the tip, it occupies the entire dome in the *etl* mutant. The localisation of RAC1 in the tip of apically growing brown algal cells has already been reported in the *Fucus* zygote rhizoid (Fowler et al., 2004) where inhibition of RAC1 by a drug resulted in a significant decrease in growth rate (Hable et al., 2008). Therefore, ETL would control the localisation of EsRAC1 to the plasma membrane of the apical cell tip, where it would ensure growth. In an *etl* mutant context, EsRAC1 would be delocalised, resulting in an inactive form that impairs polarised growth until growth arrest. By showing that EsRAC1 membrane localisation is lost in a Rho-GAP mutant, we identified a first positive regulator of EsRAC1. Interestingly, this mode of regulation by Rho-GAP is not usual for Rac proteins. In the canonical model, Rac is addressed to the membrane through isoprenylation (Yalovsky et al., 2008). This feature is probably conserved in *Ectocarpus*, as EsRac1 has the expected C-terminal isoprenylation domain, and the necessary isoprenylation machinery is found in the *Ectocarpus* proteome. Once Rac is in the membrane, it is expected to switch from its active to its inactive form under the action of RhoGAP (promoting the inactive form) and RhoGEF (promoting the active form) (see for example Hodge and Ridley, 2016). Like RhoGAPs, RhoGEFs are present in *Ectocarpus*, represented by 4 proteins with an identified Rho-GEF domain (interpro IPR000219; Uniprot: D7FPX5, D8LKY8, D7FSM5 and D7FJI7). EsRac1 can therefore undergo the expected reactions as in the canonical model. However, in this model, the Rho-GDI protein extracts the inactive Rac from the membrane and prevents its reactivation. In the pollen tube, Rho-GDI could act as a shuttle, capturing inactive GDP-bound Rac in the shanks, and delivering it to the apex where it could be reactivated by Rho-GEF (Sun et al., 2015). In *Ectocarpus*, no RhoGDI can be found in the proteome. Thus, either another factor of unknown identity plays the role of RhoGDI, or the Rac activation-inactivation switch results from a completely new mechanism. This hypothesis is supported by the fact that RhoGAP domains exhibit a surprisingly wide range of activities, often in relation to other domains of the proteins in which they are found (Amin et al., 2016). It has been observed that the BAR domain could serve as a conditional auto-inhibitor of the GAP activity (Eberth et al., 2008).

### Rac regulation of F-actin structures and vesicle trafficking

In both *etl* and WT, we observed an accumulation of actin in the most distal part of the dome, and FAs in bundles aligned along the main axis of the cell. This result is surprising, as the Rho-GTPases Rac, whose localisation is altered in *etl*, are usually involved in actin filament formation and dynamics (Rottner et al., 2017) and might be involved in cytokinesis (Moon and Zheng, 2003). In the Fucales *Silvetia compressa*, a drug against Rac1 has been shown to disrupt AFs (Muzzy and Hable, 2013). However, *de novo* nucleation of AFs occurs in some cellular structures such as filipodia protrusion in animal cells independently of Rho-GTPase (Rottner et al., 2017). In *Ectocarpus*, a role for EsRAC1 in actin filament dynamics could involve a PAK protein, which exists in the proteome as a single gene (whereas animals have multiple PAK genes, e.g. at least 6 in humans). Similarly, *Ectocarpus* has two actin depolymerisation factors, but no homolog of RIC4, a plant-specific Rac effector. As *Ectocarpus* has only one Rac protein, one would expect that a function as essential as actin (de)polymerisation would rely on this pathway, but this is not the case, at least in an *etl* context.

An abundant Golgi apparatus has previously been reported in *etl* (Le Bail et al., 2011) and it is tempting to link this apparently higher activity in vesicle trafficking to the overall greater cell wall thickness that has since been confirmed in the apical cell (Rabillé et al., 2019a). It remains unclear whether this is a direct consequence of EsRAC1 delocalisation or an indirect effect of a lower growth rate reported in the *etl* apical cell (Nehr et al., 2011). However, Hable et al. (2008) showed that Rac1 inhibition leads to delocalisation of membrane vesicles and a weaker endomembrane cycle in the zygote of the brown alga *Silvetia compressa*. Therefore, despite the significant difference between a zygote and an apical vegetative cell (Katsaros et al., 2006), both cell types confirm that Rac1 is involved in the control of vesicle trafficking in brown algae, independently of actin control. This is similar to what has been shown in yeast for the Rho3 GTPase, which regulates exocytosis in an actin-independent manner (Adamo et al., 1999), but contrasts with plants, where clathrin-mediated endocytosis is regulated by the ROP6 GTPase, in close relationship with the actin cytoskeleton (Chen et al., 2012).

### Conclusions

In summary, our results revealed that despite the conservation of key features, the regulation and function of Rac in *Ectocarpus* is far from being a copy/paste of what is already known in animals, plants or fungi, and even in the more closely related oomycetes. By studying a mutant, we have demonstrated a functional role for a BAR-Rho-GAP domain protein in the polarised growth of *Ectocarpus*. This protein appears to promote the localisation of EsRAC1 to the plasma membrane. It remains to be explored whether this localisation is correlated with the active/inactive state of EsRAC1, or whether it involves other players. Furthermore, among the cellular processes targeted by Rac, we have highlighted vesicle trafficking, with a process that does not rely on F-actin structures.

This work highlights the originality of the brown algal Rho-GTPase signalling pathway, opens the field of research on its role and regulation, and provides the first pieces of the puzzle to be assembled. The recent development of genome editing in *Ectocarpus* (Badis et al., 2021) will boost progresses.

## Methods

### Algal material

The phenotype of the mutant *etoile* (CCAP 1310/337) was generated by UV irradiation of male gametes of Ec32 (CCAP 1310/4; origin San Juan de Marcona, Peru) as indicated in Le Bail et al. (2011). In order to clear the genome from non causal mutations, *etoile* was crossed 3 times with the WT female strain (Ec568). Progeny of the F1 was screened for the [*etl*] phenotype from which female [*etl*] descendant was selected and back-crossed with the WT male Ec32 (CCAP 1310/4; origin San Juan de Marcona, Peru). A female [*etl*] descendant of this second cross was used in this study for the transcriptomic analysis. Thalli were grown in half-strength Provasoli-enriched (Starr and Zeikus, 1993) autoclaved natural seawater (pH 7.8) in a culture cabinet at 13°C with a 14:10 light: dark cycle (light intensity 29 mmol photon·m^-2^·s^-1^) as described in Le Bail and Charrier (2013). Mechanical stress consisted in placing the Petri dishes containing the algae on a rotating shaker (60 rpm) for 6 h. Cell wall digestion treatment used a cocktail of enzymes used to prepare protoplast (Benet et al., 1997).

### Identification of the mutation etoile

The mutant *etoile* (Le Bail et al., 2011) (genetic background Ec32, CCAP1310/04; male) was crossed with the strain Ec568 (CCAP1310/334; female) and a progeny of 93 gametophytes grown from a single unilocular sporangium (i.e. a single meiotic event) was first analysed for the presence of a first series of the polymorphic microsatellite markers as described in Heesch et al. (2010). 150 SSR markers separated by 10-20 cM were chosen from PCR products were analysed with the software GeneMapper v3.7. Linkage groups were formed (Lod 4, maximal distance between markers 45cM). Genetic distance was calculated from the recombination rate with the Kosambi function (Kosambi, 2016). The causal locus overlapped the supercontigs (Sctg) Sctg392 and Sctg297. Gaps between Sctgs were filled by sequencing the borders of bacterial artificial chromosomes used to assemble the genome of *Ectocarpus*, corresponding to *Ectocarpus ouroboros* mutant gDNA inserted in the pBELOBAC11 vector (Cock et al., 2010). DNA was extracted as described in Rondon et al. (1999). Missing sequence and orientation of Sctgs were confirmed by Expressed Sequence Tag (EST, cloned in plasmid pDNR222) overlapping the locus. It allowed us to insert and orientate Sctg533 and Sctg777 between the two previous Sctg392 and Sctg297 (Fig 2A). The resulting organisation of this locus is different from that obtained from predictions by Cormier et al. (2017) and Baudry et al. (2020) (Suppl Fig 1). The region was then refined by designing additional polymorphic markers using the software Websat (Martins et al., 2009) tested on 11 recombinants from a population of 688 descendants of a second Ec568 x *etl* cross. In this locus of 400 kb, we identified 15 candidate mutations from a comparison between the WT genome (Cock et al., 2010) and the *etl* genome sequenced by HiSeq2 NGS (72nt reads) using SOAP (Li et al., 2008). Each candidate mutation was sequenced from cloned PCR products amplified by proof-reading Taq polymerase. Only one was confirmed. The whole sequence of the gene *ETOILE* was assembled after Sanger sequencing of the several zones made of unidentified nucleotides (Ns). Exons were identified from a comparison between the genomic sequence and cDNAs and EST sequences and RNA-seq libraries (Suppl Table 1). The complete organisation of the gene *etoile* is shown in Fig 2B.

### Sequence analysis

The ETOILE protein sequence domains were identified using the interproscan tool (Finn et al., 2017) on the InterPro website (https://www.ebi.ac.uk/interpro/). The protein sequences were downloaded from the UniProt database (https://www.uniprot.org). Other species containing either an AH-BAR (IPR027267), I-BAR (IPR013606), or F-BAR (IPR031160) domain were chosen from UniProt (https://www.uniprot.org). The BAR domains of ETOILE and the other proteins were identified using hmmscan from the hmmer package (Eddy, 2011), then aligned with MUSCLE (Edgar, 2004). The phylogeny was reconstructed by PhyML (Guindon and Gascuel, 2003), and displayed with NJPlot (Perrière and Gouy, 1996). For rho-GTPase identification, the profiles were obtained from the Prosite database (https://prosite.expasy.org) and searched in sequences using the programs provided on this website (Sigrist et al., 2002). The sequences were aligned using MUSCLE (Edgar, 2004). The tree of Rho family protein was computed on the NGPhylogeny.fr website (Lemoine et al., 2019) using MAFFT (Katoh and Standley, 2013) for multiple alignment, BMGE (Criscuolo and Gribaldo, 2010) for alignment cleaning, FastME (Lefort et al., 2015) for phylogeny reconstruction, and iTOL (Letunic and Bork, 2021) for tree rendering.

### Transcriptomics

Laser capture experiments were performed as described in Billoud et al. (2021). In short, *Ectocarpus* sporophytes were grown directly on PEN slides until they reached a stage of ∼ 30 cells. Then they were fixed in acetone and the different cell types were captured by laser microdissection (Carl Zeiss PALM MicroBeam unit with the PalmRobo 4.5 software) as described in detail in Saint-Marcoux et al. (2015). RNAs were extracted using the Arcturus PicoPure RNA extraction kit and amplified in triplicates with the Ovation RNA-seq System v2 kit from NuGEN. Amplified cDNAs were either sequenced, or amplified by Q-PCR. For each cell type 10 Millions of Paired-End reads were produced using the HiSeq NGS technology, and analysed as described in detail in Billoud et al. (2021). Briefly, reads were filtered and mapped onto the *Ectocarpus* genome and transcriptome V2 representing 18271 genes (Cormier et al., 2017) available in Orcae (Sterck et al., 2012). Differential Gene Expression was detected using edgeR (McCarthy et al., 2012; Robinson et al., 2010) setting the maximum False Discovery Rate (FDR) to 5 × 10^−2^. In parallel, cDNAs were amplified by Q-PCR using 5’ and 3’ oligonucleotides (Eurogentec) specific to each gene. Oligonucleotides were designed with the software Perlprimer (Marshall, 2004). List of oligonucleotides is given in Suppl Table 2. cDNA amplification and quantification took place as described in Billoud et al. (2021).

### Over-expression of the ETOILE BAR and ETOILE RHO-GAP protein domains

Sequences were analysed to detect potential signal peptide and transmembrane domain using SignalP-5.0 (Almagro Armenteros et al., 2019) and TMHMM 2.0 (Krogh et al., 2001) online applications. In order to delineate each module, BlastP queries were performed using the UniProt and the Protein Data Banks. The precise delineation of each module was refined by performing Hydrophobic Cluster Analysis plots (Callebaut et al., 1997) Specific forward and reverse primers (Suppl Table 3) were designed using the In-Fusion Cloning Primer Design Tool from TaKaRa (https://www.takarabio.com/learning-centers/cloning/primer-design-and-other-tools) and used to amplify the BAR (residues 103 to 374) and the RHO GAP (Residues 376 to 579) domain sequences (Suppl Fig 2). The purified PCR products were cloned into the expression vector pFO4 predigested with EcoRI and BamHI using the In-Fusion HD Cloning kit (TaKaRa). This plasmid, resulting from a modification of the pET15b plasmid (Novagen, USA) bears a sequence encoding an hexa histidine tag upstream from the multi cloning site (Groisillier et al., 2010). Recombinant plasmids in *E. coli* Stellar chemically competent cells (TaKaRa) were sequenced prior to transformation of the BL21(DE3) competent *E. coli* (BioLabs) strains. Unless otherwise stated, all other chemicals were from Sigma.

### Production and purification of BAR and RHO-GAP modules

Unless otherwise stated, experiments were performed at 4°C and the strategy used was the same for both modules. Purification steps were performed using an AKTA Avant apparatus (Cytiva). Recombinant *E. coli* BL21(DE3) strains were grown for 72 hours at 20°C in 1L ZYP medium (Studier, 2005) containing 100 µg.mL^-1^ ampicillin. Cultures were stopped by centrifugation at 3000g for 20 min. The pellets were kept at -20°C until further used. Pellets corresponding to ∼200 mL culture medium were suspended in 6mL of 10 mM TrisHCl (pH8.0), 25% sucrose and 2 mg lysozyme. The sample was stirred for 15 minutes prior to the addition of 12 mL of 20 mM TrisHCl, 1% deoxycholate, 1% Triton X-100 and 100 mM NaCl (pH 7.5). The sample was further incubated for 5 min at 4°C. 20 µL of DNAse I (500 Units.µL^-1^) and MgCl2 at 5 mM final concentration were added. The sample was stirred until the viscosity dropped. After centrifugation for 1 hour at 29000 g, the cell free supernatant was further filtrated on 0.2 µm cellulose acetate and diluted three times in 50 mM TrisHCl, 100 mM NaCl and 5 mM MgCl2 (pH7.8; Buffer A) before being loaded at a flow rate of 1 mL.min^-1^ on top of a 5 mL Histrap FF column. The column was washed at the same flow rate with this buffer until the absorbance at 280 nm was less than 40mAU. A subsequent wash was carried out at 2 mL.min^-1^ with 2.5 % buffer B (same as buffer A but containing 1 M imidazole). When the absorbance at 280 nm was less than 25 mAU, the elution of proteins was performed using a linear gradient from 2.5% B to 100% buffer B in 22 Column volumes (CV). 2 mL fractions were collected during the elution step and analysed by SDS PAGE. 2 µL of each fractions were also spotted on nitrocellulose membrane to perform Dot Blot. Monoclonal anti-polyhistidine peroxidase antibodies (Sigma) were used at a final concentration of 1/10000 to specifically detect the his-tagged protein. Revelation was performed by chemiluminescence using the Clarity Western Substrate (BioRad) and visualization was carried out using the Chemi-Capt software. Fractions containing the pure BAR module were pooled and dialysed against 3x 1L 50 mM Tris HCl + 100 mM NaCl (Buffer C). They were concentrated using Amicon-Ultra 15 or 0.5 (10 kDa) devices. Because the Rho-GAP module was not expressed as well as the BAR module, pellets corresponding to ∼1 L culture were used and downstream buffer and chemical volumes increased accordingly. Two 5 mL HisTrap FF columns (Cytiva) in series were used with washing and elution steps performed as described above. Fractions containing the his-tagged protein were analysed by SDS-PAGE and Western blot. Transfer from SDS PAGE onto ready to use 0.2 µm nitrocellulose membrane (BioRad) was performed in 7 min at 1.3 A and 25 V using a Trans-Blot TurboTM transfer system (BioRad). Recognition and revelation of the his-tagged protein was performed as described for the Dot Blot. All the fractions containing the tagged protein were pooled and concentrated on Amicon Ultra devices prior to being loaded on top of a Superdex200 Increase 10/300 GL (Cytiva) previously equilibrated in Buffer C. Protein elution was performed at a flow rate of 0.5 mL.min^-1^ and 0.5 mL fractions were collected. Purity of the RHO-GAP module was further checked by SDS PAGE analysis.

Protein amount in desalted samples was estimated at 280 nm using a ThermoS Scientific NanoDrop One Spectrophotometer. The molar extinction and molecular weight of both BAR and RHO-GAP modules were deduced from the amino acid sequence using the ProtParam program (Gasteiger et al., 2005). BAR module molar extinction coefficient and molecular weight were 13,200 M^-1^cm^-1^ and 30.7 kDa respectively. The molar extinction coefficient of the RHO-GAP module was estimated at 16,180 M^-1^cm^-1^ and its molecular weight was valued at 23.810 kDa.

### Production of specific ETOILE-BAR and ETOILE-RHO-GAP antibodies

Antibodies against the ETOILE BAR and RHO-GAP domains were produced in rabbits (Covalab: https://www.covalab.com/). ELISA titration of the serum and purified antibodies raised against BAR were > 64000 and < 8 ng.mL^-1^ respectively, and against RHO-GAP 16000 for the Serum and 16 ng.mL^-1^ respectively. They were stored in 50% 0.1 M Tris / Glycine pH 7.8, 0.02% azide de sodium and 50% glycerol -20°C for several months. Sera collected at day 0 showed no immunoreactivity with the two expressed protein domains (shown by western blot).

### Western Blot with total proteins

*Ectocarpus* filaments were ground in liquid nitrogen and ∼ 1 mg of fine powder was added to 1.5 mL of extraction buffer (6M Urea, 2M Thiourea, 1% DTT, 4% CHAPS) in a 50 mL Falcon tube. After shaking (vortex) at 4°C for 20 min, the tube was centrifuged at 13,000g for 30 min at 40°C. Proteins in the supernatant were used immediately or aliquoted and stored at -80°C in 50% glycerol. Purified proteins were mixed with 1X sample buffer (60mM Tris-HCl pH6.8; 7.5% glycerol, 1.5% SDS, 1.5% DTT, 0.02% bromophenol blue and heated for 5 min at 94°C. They were run on a Gel SDS-PAGE 4-20% acrylamide and stained with Coomassie blue for 30 min, followed by destaining in 30%:10% Ethanol:acetic acid for variable durations. Western blot was carried out as described above.

### Immunolocalisation

This protocol is adapted from other immunolocalisation protocols (Katsaros, 1992; Le Bail et al., 2010). RAC1, Rho-GAP and BAR antibodies: *Ectocarpus* Ec32, *etl* and U749 (progeny of Ec568 x *etl*) were grown on a glass slide or coverslip until they reach ∼ 100 cells in sterile sea water (SW). Slide was drained from SW, put in a dry clean Petri dish (PD) and fixed with 600 µL of 3 % paraformaldehyde (PFA) in SW for 3 h at 4°C. After rinsing the slide 3x 10 min with a large volume of PBS (137 mM NaCl, 2.7 mM KCl, 10 mM Na2HPO4, 1.8 mM KH2PO4, pH7.4) - 0.5 % Triton, it was transferred onto a piece of hydrophobic parafilm laid on a damp Wattman paper (PBS) in a new PD. ∼ 600 µL of cell wall degradation solution (2 % cellulase, 2 % hemicellulase, 1 % driselase, 1 % macerozyme, 0.5 % pectinase, 1 to 2 % alginate lyase, 0.2 % Triton) were added, and the slide incubated at room temperature (RT) for 20min in the humid chamber. After 4x 10min abundant washes with PBS-Triton, ∼ 600 µL of blocking solution (3 % BSA in 0.5 % Triton X100-PBS) were added and incubated for 1 hour at RT (or O/N at 4°C). After rinsing twice 10min with 1 % BSA in PBS, 100 µL of polyclonal antibodies ETOILE-BAR and ETOILE-RHOGAP (1:500), or of the rabbit anti-Human Rac1 polyclonal antibodies (1:40, Thermofisher PA1-091) were added in 1 % BSA-PBS and incubated O/N at 4°C. After incubation 1x 10 min in a large volume of PBS, 1x 10 min in 1% BSA-PBS and 1x 10min in 500 µL 1% BSA-PBS, the slides were transferred back to the humid chamber on a clean piece of parafilm. 150 µL of 1% BSA-PBS containing 1:200 anti-rabbit FITC-IgG secondary antibody (Sigma, F6258) were added and incubated 1-2 h at RT in the dark. After rinsing 1x 1min in 1 mL (enough to cover the surface with filaments) 1% BSA-PBS, 1×10 min in 1 mL 1% BSA-PBS and 4x 8 min in 2-3 mL PBS, the slides were mounted in Vectashield (Vector Laboratories), sealed and observed with an HCX PL APO lambda blue 63X 1.4 oil immersion objective mounted on a SP5 confocal microscope (Leica) to detect an FITC signal (Ex 495 nm; Em 525 nm). Same laser power and pinhole were used for negative controls (with only the secondary antibody) and positive samples.

### Actin antibody

Indirect immunofluorescence (IIF) assay was performed as described earlier (Mermelstein et al., 1998) with a slight modification in the blocking conditions, that was carried out at 4°C overnight.. Primary antibody was rabbit anti-plant actin (AS13 2640, Agrisera; dilution 1:300) and second antibody goat anti-rabbit IgG conjugated to Alexa Fluor 488 (Thermo Fisher Scientific, A-11008; dilution 1:200).

### Labelling of actin filaments

This actin labelling protocol is a modified version of the one in Rabillé et al. (2018). All the steps took place at RT. The material was prefixed in a prefixation solution containing 0.20% Triton X-100, 2% DMSO and 300 μM 3-maleimidobenzoic acid N-hydroxysuccinimide ester (MBS; Sigma) dissolved in microtubule stabilization buffer specific for algae (MTB; 50 mM PIPES, 5 mM MgSO4, 5 mM EGTA, 25 mM KCl, 4 % (w/v) NaCl, 2.5 % (w/v) PVP, 1 mM DTT, pH 7.4). The material was incubated for 30 min in the dark and then washed with MTB once. For fixation, the material was incubated for 40 min in the dark in a fixation solution containing 2% PFA. After the incubation the material was washed in gradients of MTB:PBS solution (Phosphate buffer solution; 137 mM NaCl, 0.7 mM KCl, 5.1 mM Na2HPO4, 1.7 mM KH2PO4, 0.01% NaN3, pH 7.4). After the washes, the material was incubated for 20 min in a solution of 2% (w:v) Onozuka R-10 (Yakult Honsha Co., Tokyo, Japan), 2% Hemicellulase (Sigma), 1% driselase (Sigma), 0.5% pectinase (Sigma) 1 % macerozyme R-10 (Yakult), 50 U.mL^-1^ Alginate lyases -G (isolated from *Zobellia galactonivorans* (Rabillé et al., 2019b)), dissolved in 1:1 MTB:PBS containing 0.12 % Triton X-100 and 0.17 μM Phalloidin-Alexafluor 488 conjugate (Invitrogen). After washes with MTB:PBS 1:1, an extraction step took place with a solution of 5% DMSO and 3% Triton X-100 in MTB:PBS 1:1. After washes, the filaments were incubated in the actin labeling solution containing 0.33 μM Phalloidin-Alexafluor 488 conjugate and 0.1 % Triton X-100 in 1:1 MTB:PBS for 1 h in RT. Finally, the material was washed with PBS and then mounted in Vectashield Vibrance (Vectorlabs).

## Acknowledgement

We are grateful to Élodie Rolland for providing *Ectocarpus* Ec32 material and maintaining the *etl* progeny in stock, Mark Cock for making the two BAC clones available, and Lionel Cladière for his help with the design of the expressed protein domains. The European COST Action FA1406 “Phycomorph” is acknowledged for funding scientific exchange missions of Z. Nehr, A. Nasir and H. Rabillé.

Z. Nehr was funded by CNRS PhD program and H. Rabillé by Region Bretagne and Sorbonne University (project “Ectotip”).

## Notes

### Competing Interest Statement

The authors have declared no competing interest.

