## Supplementary Figure 1 for "Tip growth in the brown alga *Ectocarpus* is controlled by a RHO-GAP-BAR domain protein independently from F-actin organisation"

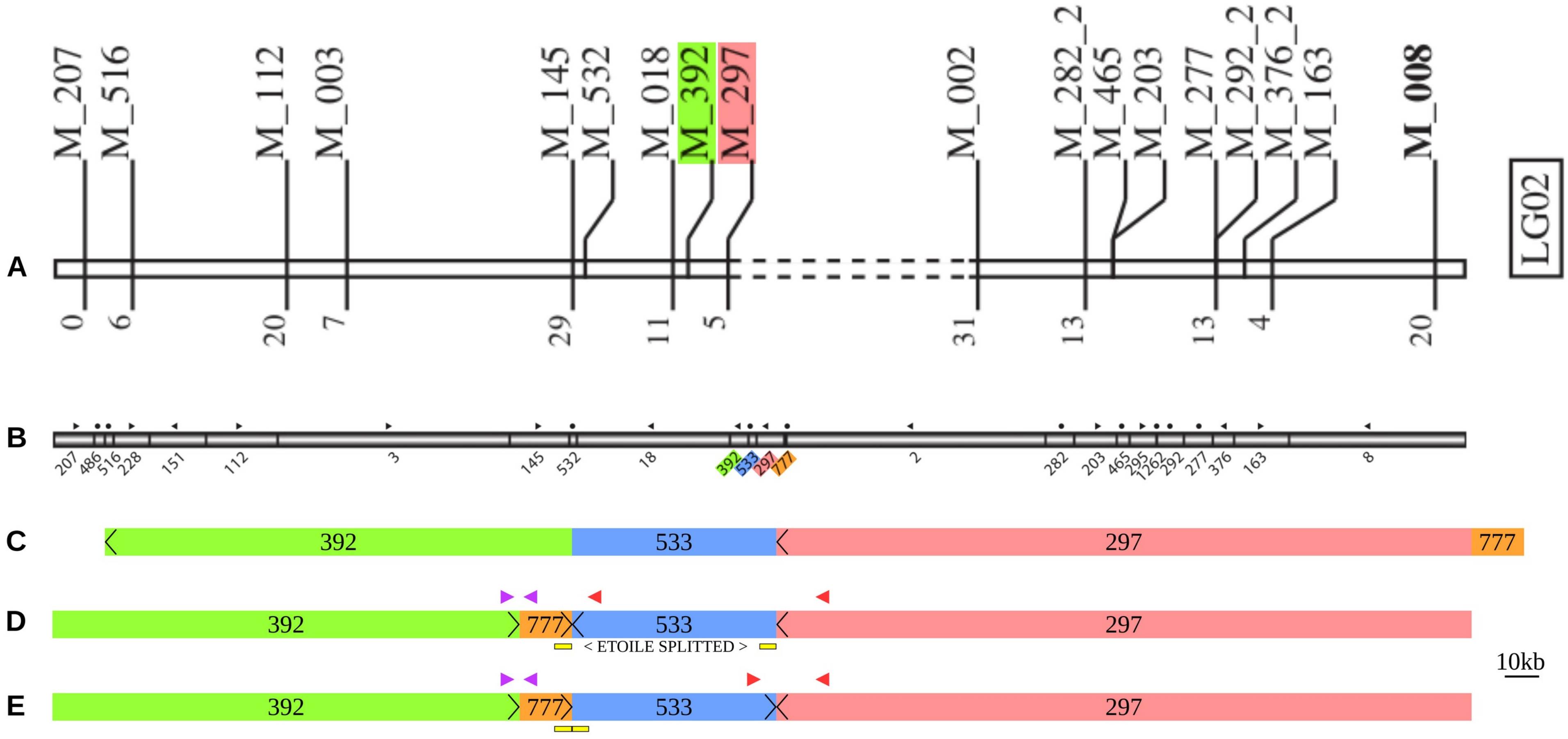


*Suppl figure 1 : Assembly of the super-contigs in the pseudo chromosomes.* A: genetic map assigning super-contigs 392 and 297 to the linkage group 02 (Heesch et al. 2010). B and C: Enhanced map adding super-contigs 533 and 777, without orientation, and details of the region containing the locus *etoile* as it appears in the V2 of the genome (Cormier et al. 2017). D: The region after re-assembly (Baudry 2020) Purple and red arrowheads show the positions of two BAC-ends. The position and orientation of sctg_392 and 777 are in agreement with the purple one. Sctg_533 is mis-oriented, as shown by the position of the red BAC-ends. As a result, the gene ETOILE (yellow) is splitted into two parts. E: Correct order and orientation of the super-contigs, in which ge gene ETOILE is found as one contiguous sequence (Nehr et al 2011).
