## Supplementary Figure 2 for "Tip growth in the brown alga *Ectocarpus* is controlled by a RHO-GAP-BAR domain protein independently from F-actin organisation"

```
        10         20         30         40         50         60
MRRKQQSSSG AGAVSNHDLY DTDPPRPGRL LGMPAPVPAP QGGAAAFNAA GRGMSFPGAT

        70         80         90        100        110        120
AASGGGQGGG GGSGGGGAGG GGGGVDLDLR LRDSSQKGGK RGGGFRDWKR RVGEKAILKI

       130        140        150        160        170        180
GLAAPSNDAN TGDIEQPLHS FVQMRDGLKD IQAQVRNYVL AMEGLFGAAH NMTFLFAPLS

       190        200        210        220        230        240
GETAKQLKLS LCRVAVQDAM IRGLTEYLEA PLKAVLDFCD GIDLLAKRRG QLLLDYGHHK

       250        260        270        280        290        300
RKHEAVEARR ESGKGGDMEE EVKARSTKLN ASIDNLKDCT TQILAQVAAL SKVEPQLKES

       310        320        330        340        350        360
LFKGLSACQV FYCQASLSAF ETDFVESTDM RDYVMNLQST TTTLENSMPL PPLGEHVAFL

       370        380        390        400        410        420
RNALASAAAE VRGEGKLGYV YGRPLSDLEE PPPLLLECIG FLDAHGLYVP GLFRVPGNTE

       430        440        450        460        470        480
KIQQLKSTFD AGEHVEFDER IHDVHEVATL LKMYFRELPE PVVPTAFYYP MLEASVKDLT

       490        500        510        520        530        540
EFVETVRVLV AQLPEINRTC VRLLFAFLLR VSLLSAENLM TTENLGIVFA PNLLRAPNAT

       550        560        570
EMNVMDLQAC INLVKRIIEC AKEIFPGLWQ GYPVVASQG
```

*Supplementary figure 2: The two domains BAR and RHO-GAP of the protein ETOILE were overexpressed in vitro.* The BAR module (underline in yellow) and the RHO-GAP module (blue) were overexpressed as desribed in the Method section. Starting position and ending position as predicted by Interproscan are indicated, with plain lines for the most restrictives and dashed lines for the most permissive positions for each domain (BAR : Starting position : between 103 and 136 AA; Ending site : 321-342 ; RHO-GAP : Starting position : between 355 and 398 AA ; Ending site : 539-574). BAR module is 29.94 kDa and RHO-GAP module is 22.99 kDa.
