## Supplementary Figure 3 for "Tip growth in the brown alga *Ectocarpus* is controlled by a RHO-GAP-BAR domain protein independently from F-actin organisation"

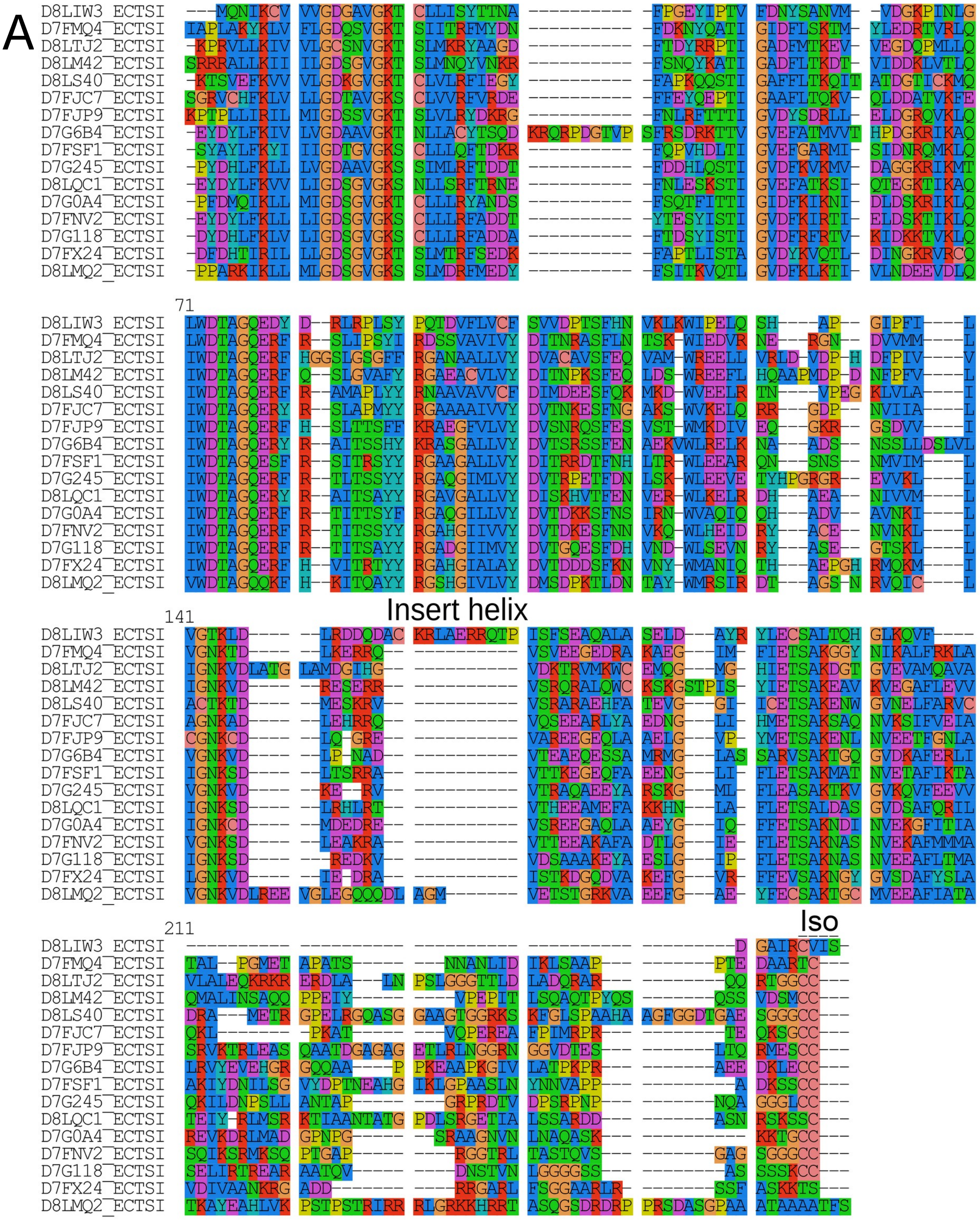
A
71
141
Insert helix
211
Iso

B

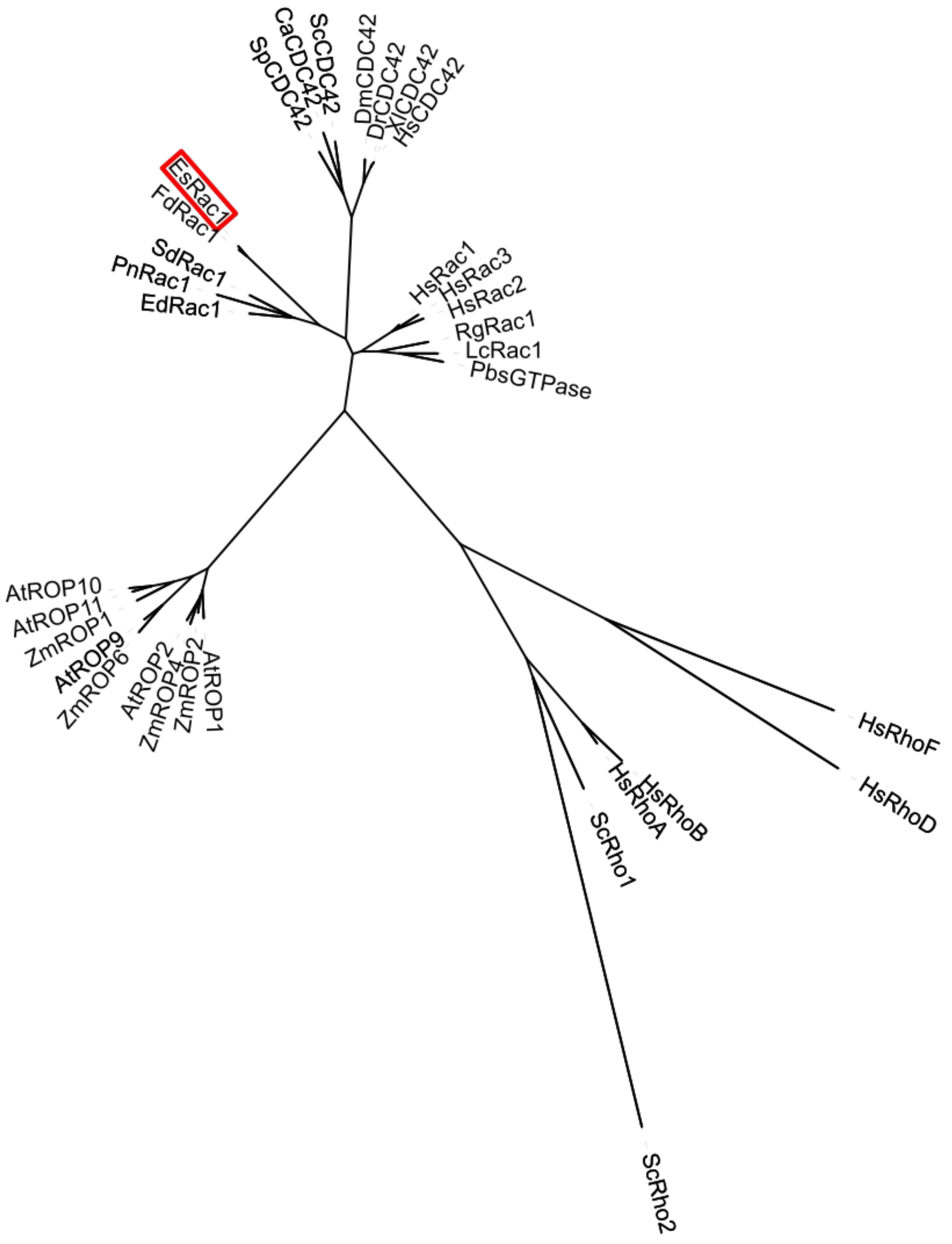


*Supplementary figure 3: characterisation of EsRAC1* A: Multiple alignment of the characteristic domains of all small GTPases in *Ectocarpus*. The only Rho-GTPase is D8LIW3, which can be identified by the specific insert helix and the isoprenylation site (marked above the alignment). B: Phylogenetic tree of Rho-GTPase sub-families. EsRAC1 is framed in red. Species abbreviations: At: *Arabidopsis thaliana*, Ca: *Candida albicans*, Dm: *Drosophila melanogaster*, Dr: *Dario rerio*, Ed: *Eurychasma dicksonii*, Hs: *Homo sapiens*, Lc: *Lichtheimia corymbifera*, Pb: *Phycomyces blakesleeanus*, Pn *Phytophthora nicotianae*, Rg: *Rhodotorula graminis*, Sd: *Saprolegnia diclina*, Sp: *Saccharomyces scerevisiae*, Xl: *Xenopus laevis*, Zm: *Zea mays*.

References of the sequences used:

| Name | Unipro accessiont |
|---|---|
| AtROP1 | P92978 |
| AtROP2 | Q38919 |
| AtROP9 | O82480 |
| AtROP10 | Q9SU67 |
| AtROP11 | O82481 |
| CaCDC42 | P0CY33 |
| DmCDC42 | P40793 |
| DrCDC42 | Q6TH34 |
| EdRac1 | A0A1B3SNZ7 |
| EsRac1 | D8LIW3 |
| FdRac1 | Q6T388 |
| HsCDC42 | P60953 |
| HsRac1 | P63000 |
| HsRac2 | P15153 |
| HsRac3 | P60763 |
| HsRhoA | P61586 |
| HsRhoB | P62745 |
| HsRhoD | O00212 |
| HsRhoF | Q9HBH0 |
| LcRac1 | A0A068S7E8 |
| PbsGTPase | A0A167RDT6 |
| PnRac1 | A0A0W8CIX0 |
| RgRac1 | A0A194SDH4 |
| ScCDC42 | P19073 |
| ScRho1 | P06780 |
| ScRho2 | P06781 |
| SdRac1 | T0RB17 |
| SpCDC42 | Q01112 |
| XlCDC42 | A0A1L8FGA0 |
| ZmROP1 | E9P1P1 |
| ZmROP2 | Q9XF06 |
| ZmROP4 | Q9XF08 |
| ZmROP6 | Q9LEC5 |
