## Supplementary Figure 4 for "Tip growth in the brown alga *Ectocarpus* is controlled by a RHO-GAP-BAR domain protein independently from F-actin organisation"

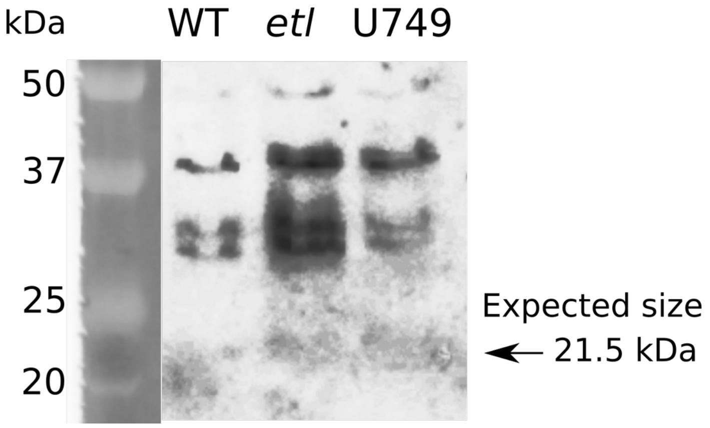


*Supplementary figure 4. Western blot of total Ectocarpus proteins labelled with HsRac1 polyclonal antibody.*
