## Supplementary Table 3 for "Tip growth in the brown alga *Ectocarpus* is controlled by a RHO-GAP-BAR domain protein independently from F-actin organisation"

*Supplementary table 3: Synthetic oligonucleotides used to amplify by PCR the BAR and RHO modules***.** The restriction site sequences are underlined and written in bold. The sequences specific of the plasmid are in italic.

| Module | Primer Forward | Primer reverse |
|---|---|---|
| **BAR** | *5'-CCATCACCAT***GGATCC**GGCGGCTTCAGGGACTGG | 5'-*CAGCCGGATC***GAATTC**TCACTCCCCGCGAACCTC |
| **RHO-GAP** | 5'-*CCATCACCAT***GGATCC**AAGCTGGGGTATGTCTACGGC | 5'-*CAGCCGGATC***GAATTC**TCAACCCTGCGAGGCCACC |
