## Supplementary Table 4 for "Tip growth in the brown alga *Ectocarpus* is controlled by a RHO-GAP-BAR domain protein independently from F-actin organisation"

*Supplementary Table 4: Occurrences of Prosite GTPase profiles in Ectocarpus predicted GTPases.* For each protein, the profile name, raw score, adjusted score and position within the sequence are shown, each with a color: pink for Rho-GTPAse, blue, for Rab GTPase, yellow for Ras GTPase (ordered by best score). Protein Accession number (AC), identity (ID) and function (as they appear in Uniprot) are colored like the profile for which they score the best.

| Profile | Raw | Score | beg | end | AC | ID | Function |
|---|---|---|---|---|---|---|---|
| PS51420 | 7537 | 26.573 | 1 | 176 | D8LIW3 | D8LIW3_ECTSI | RAC, RHO family GTPase GN=RAC |
| PS51419 | 4154 | 19.856 | 1 | 196 | | | |
| PS51421 | 3223 | 16.473 | 1 | 196 | | | |
| PS51419 | 7168 | 33.062 | 3 | 195 | D7FJC7 | D7FJC7_ECTSI | Rab5, RAB family GTPase GN=Rab5 |
| PS51421 | 3566 | 18.087 | 2 | 195 | | | |
| PS51420 | 3276 | 9.274 | 2 | 172 | | | |
| PS51419 | 6594 | 30.547 | 14 | 222 | D7FJP9 | D7FJP9_ECTSI | Rab8D, RAB family GTPase GN=Rab8D |
| PS51421 | 3621 | 18.346 | 14 | 223 | | | |
| PS51420 | 3838 | 11.556 | 13 | 181 | | | |
| PS51419 | 6876 | 31.782 | 16 | 215 | D7FMQ4 | D7FMQ4_ECTSI | Rab6, RAB family GTPase GN=Rab6 |
| PS51421 | 3424 | 17.419 | 17 | 198 | | | |
| PS51420 | 3480 | 10.102 | 15 | 188 | | | |
| PS51419 | 8596 | 39.319 | 4 | 203 | D7FNV2 | D7FNV2_ECTSI | Rab1B, RAB family GTPase GN=Rab1B |
| PS51421 | 4207 | 21.104 | 1 | 203 | | | |
| PS51420 | 3738 | 11.15 | 2 | 171 | | | |
| PS51419 | 7568 | 34.814 | 2 | 208 | D7FSF1 | D7FSF1_ECTSI | Rab2A, RAB family GTPase GN=Rab2A |
| PS51421 | 3386 | 17.24 | 1 | 172 | | | |
| PS51420 | 3162 | 8.811 | 1 | 168 | | | |
| PS51419 | 6567 | 30.428 | 133 | 333 | D7FX24 | D7FX24_ECTSI | Rab8C, RAB family GTPase GN=Rab8C |
| PS51421 | 2714 | 14.078 | 125 | 336 | | | |
| PS51420 | 3140 | 8.722 | 131 | 301 | | | |
| PS51419 | 7714 | 35.454 | 7 | 205 | D7G0A4 | D7G0A4_ECTSI | Rab8A, RAB family GTPase GN=Rab8A |
| PS51421 | 4512 | 22.539 | 4 | 205 | | | |
| PS51420 | 3590 | 10.549 | 5 | 188 | | | |
| PS51419 | 8066 | 36.996 | 5 | 202 | D7G118 | D7G118_ECTSI | Rab1A, RAB family GTPase GN=Rab1A |
| PS51421 | 3791 | 19.146 | 1 | 202 | | | |
| PS51420 | 3452 | 9.989 | 3 | 175 | | | |
| PS51419 | 7479 | 34.424 | 8 | 209 | D7G245 | D7G245_ECTSI | Rab18, RAB family GTPase GN=Rab18 |
| PS51421 | 3108 | 15.932 | 4 | 210 | | | |
| PS51420 | 3574 | 10.484 | 6 | 175 | | | |
| PS51419 | 5757 | 26.879 | 8 | 229 | D7G6B4 | D7G6B4_ECTSI | Rab11B, RAB family GTPase GN=Rab11B |
| PS51421 | 2344 | 12.337 | 5 | 214 | | | |
| PS51420 | 3467 | 10.05 | 6 | 191 | | | |
| PS51419 | 6969 | 32.19 | 3 | 213 | D8LM42 | D8LM42_ECTSI | Rab7, RAB family GTPase GN=Rab7 |
| PS51421 | 3832 | 19.339 | 3 | 212 | | | |
| PS51420 | 3507 | 10.212 | 2 | 181 | | | |
| PS51419 | 5966 | 27.795 | 49 | 282 | D8LMQ2 | D8LMQ2_ECTSI | Rab8E, RAB family GTPase GN=Rab8E |
| PS51421 | 3254 | 16.619 | 49 | 332 | | | |
| PS51420 | 3255 | 9.189 | 47 | 227 | | | |
| PS51419 | 7651 | 35.178 | 7 | 214 | D8LQC1 | D8LQC1_ECTSI | Rab11A, RAB family GTPase GN=Rab11A |
| PS51421 | 3290 | 16.789 | 4 | 219 | | | |
| PS51420 | 3496 | 10.167 | 5 | 173 | | | |
| PS51419 | 6044 | 28.137 | 8 | 225 | D8LS40 | D8LS40_ECTSI | Rab50, RAB family GTPase GN=Rab50 |
| PS51421 | 3056 | 15.688 | 8 | 226 | | | |
| PS51420 | 3141 | 8.726 | 6 | 180 | | | |
| PS51419 | 5390 | 25.271 | 149 | 364 | D8LTJ2 | D8LTJ2_ECTSI | Rab7L, RAB family GTPase GN=Rab7L |
| PS51421 | 2606 | 13.57 | 146 | 365 | | | |
| PS51420 | 3221 | 9.051 | 147 | 325 | | | |
